# Time-lapsed proteomics reveals a role for the novel protein, SNED1, in modulating ECM composition and protein folding

**DOI:** 10.1101/2022.01.13.476092

**Authors:** Fred Lee, Xinhao Shao, Yu (Tom) Gao, Alexandra Naba

## Abstract

The extracellular matrix (ECM) is a complex and dynamic meshwork of proteins providing structural support to cells. It also provides biochemical signals governing cellular processes including proliferation and migration. Alterations of ECM structure and/or composition has been shown to lead to, or accompany, many pathological processes including cancer and fibrosis. To understand how the ECM contributes to diseases, we first need to obtain a comprehensive characterization of the ECM of tissues and of its changes during disease progression. Over the past decade, mass-spectrometry-based proteomics has become the state-of-the-art method to profile the protein composition of ECMs. However, existing methods do not fully capture the broad dynamic range of protein abundance in the ECM, nor do they permit to achieve the high coverage needed to gain finer biochemical information, including the presence of isoforms or post-translational modifications. In addition, broadly adopted proteomic methods relying on extended trypsin digestion do not provide structural information on ECM proteins, yet, gaining insights into ECM protein structure is critical to better understanding protein functions. Here, we present the optimization of a time-lapsed proteomic method using limited proteolysis of partially denatured samples and the sequential release of peptides to achieve superior sequence coverage as compared to standard ECM proteomic workflow. Exploiting the spatio-temporal resolution of this method, we further demonstrate how 3-dimensional time-lapsed peptide mapping can identify protein regions differentially susceptible to trypsin and can thus identify sites of post-translational modifications, including protein-protein interactions. We further illustrate how this approach can be leveraged to gain insight on the role of the novel ECM protein SNED1 in ECM homeostasis. We found that the expression of SNED1 by mouse embryonic fibroblasts results in the alteration of overall ECM composition and the sequence coverage of certain ECM proteins, raising the possibility that SNED1 could modify accessibility to trypsin by engaging in protein-protein interactions.

## INTRODUCTION

The extracellular matrix (ECM) is a remarkably complex assembly of proteins that plays a critical architectural role in multicellular organisms by governing cell polarization, organizing cells into tissues, conferring mechanical properties to tissues and organs, and contributing to morphogenesis [1–4]. The ECM also conveys biochemical and biomechanical signals that tightly control all cellular processes from cell proliferation and survival, to migration, to differentiation [1,5]. Alterations of ECM structure and/or composition arising from mutations in ECM genes, or from an imbalance in ECM homeostasis, *e*.*g*., excessive accumulation or degradation, lead to, or accompany, many pathological processes including skeletal diseases, cardiovascular diseases, fibrosis, and cancer [6–12]. To understand the underlying molecular bases of ECM involvement in diseases, we need methods and tools capable of probing the complexity of this compartment.

Over the past decade, bottom-up mass-spectrometry (MS)-based proteomics, has become a method of choice to profile the global protein composition of ECMs – or matrisomes – of tissues, organs, or produced by cells *in vitro* [13–17]. This method infers protein presence based on the identification of the most abundant peptides in any given sample. Proteomics has revealed that the ECM of any given tissue is made of well over 100 distinct proteins and has allowed the identification of ECM protein signatures characteristic of physiological processes (*e*.*g*., development [18], aging [19–22]) and pathological states (*e*.*g*., cardiovascular diseases [23], cancer [24], fibrosis [25–27]). However, existing proteomic approaches present limitations. First, the size and relative abundance of proteins found in the ECM span a very broad dynamic range, from very large and hyper-abundant collagens to small and low-abundance ECM remodeling enzymes or ECM-bound growth factors. As such, and despite improvements in instrumentation and sample preparation, including protein and peptide fractionation, proteins present in lower abundance are often eclipsed [14]. In addition, trypsin digestion remains the gold standard to generate peptides in a bottom-up MS workflow [28] but is not without limitations. For example, some proteins can be intrinsically resistant to tryptic digestion and, as such, may not be detected. ECM proteins are often highly cross-linked and cannot be easily solubilized, thus limiting trypsin accessibility. To overcome this, we and others previously proposed that hard-to-digest ECM proteins would benefit from sequential multi-proteases digestion (in our case LysC + trypsin) [17,29]. Last, lower abundance proteins found in the ECM (proteins we have previously grouped under the term “matrisome-associated and including ECM-affiliated proteins, ECM regulators, and ECM-bound secreted factors [17,30]) tend also to be smaller, and so generate less peptides and, consequently, are less likely to be identified. Yet, these proteins are critical regulators of ECM homeostasis and functions. We and others have contributed to the enhancement of proteomic methods, however, until now, the focus has remained on improving protein identification and increasing the number of proteins identified [14,31,32].

Moreover, like all other proteins, ECM proteins exist in multiple proteoforms arising from alternative splicing for isoforms, from single-nucleotide variants, or from post-translational modifications [33,34] and these proteoforms can play important roles in health and disease. While proving the presence of a protein in a given sample only needs a few unique peptides, capturing a specific proteoform requires the detection of unique peptide sequences that cover all the variation and modification sites of a given protein, which often necessitates higher sequence coverage. Of note, the combined average sequence coverage of ECM proteins in MatrisomeDB, a database compiling ECM proteomic datasets [35], is 36.85% with a median sequence coverage <30%. The addition of new datasets to MatrisomeDB contributes little to improving overall sequence coverage, as the same trypsin-based proteomics protocol simply repeats previous observations. This implies that conventional “identification-oriented” proteomic workflows cannot meet the need for studies of ECM proteoforms. Instead, we propose to steer the development towards “coverage-oriented” approaches to increase the sequence coverage of ECM proteins and thus obtain more information on ECM proteoforms.

Last, certain critical aspects of ECM biochemistry remain largely understudied, including folding and ECM protein-protein interactions, in the context of the assembled and insoluble ECM meshwork, as opposed to studying individual ECM proteins or fragments in soluble forms. Yet, we know that ECM protein folding and the nature of the protein complexes formed in the ECM are of paramount importance to achieve proper ECM function.

Limited proteolysis coupled to proteomics has emerged as a powerful method to elucidate protein structures and conformational changes, and to map sites of protein-protein interactions [36,37]. However, until now, this approach has mainly been used to study soluble proteins, as they provide omnidirectional access for proteases. Here, we used limited proteolysis to devise a time-lapsed digestion pipeline to study the ECM with two goals: 1) to release peptides *sequentially* over time and thus generate peptide samples of lower complexity, with the goal of enhancing protein identification and protein coverage, 2) since ECM proteins are highly insoluble and can only be partially solubilized prior to tryptic digestion [17,38] (*see also* Figure 1A), a time-lapsed tryptic digestion will result in the release of peptides from more exposed or accessible regions of proteins at earlier timepoints, and of peptides from buried protein domains as the digestion progresses. As a result, the mapping of peptides obtained via time-lapsed tryptic digestion can provide important information on protein folding and possible sites of protein-protein interactions.

**Figure 1.**
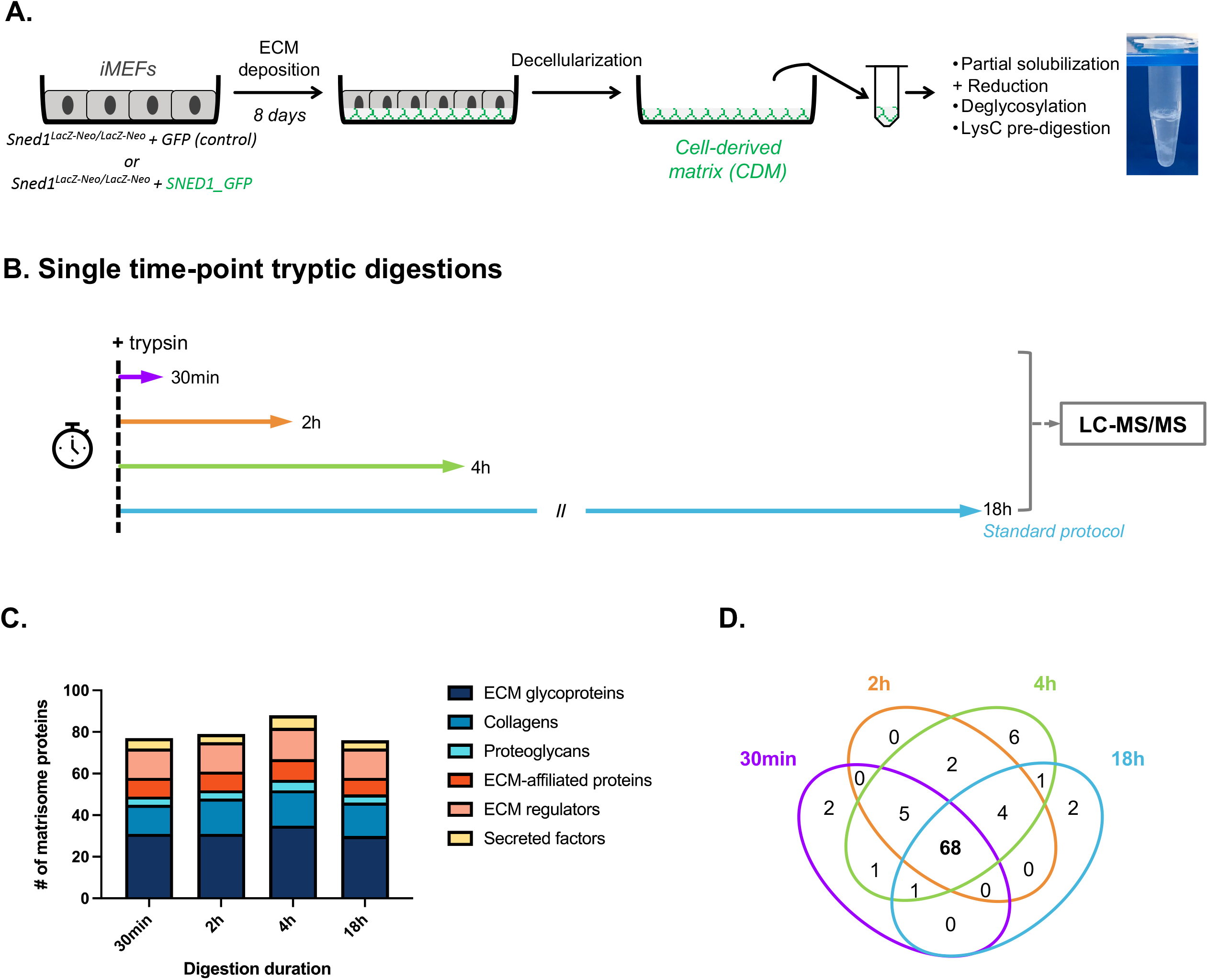
Identification of ECM proteins using time-restricted proteolysis protocol. **A**. Schematic representation of the experimental pipeline including the production of cell-derived matrix (CDM) by immortalized mouse embryonic fibroblasts (MEFs) over the course of 8 days and the initial steps of sample preparation prior to LC-MS/MS analysis. **B**. Schematic representation of the time-limited proteolysis protocol using single timepoint digestions of defined duration (30 minutes, 2 hours, 4 hours, or 18 hours). **C**. Bar graph depicts, for each matrisome category, the number of ECM proteins identified in two biological replicates (intersection), using single timepoint digestions (*see also Supplementary Table 2*). **D**. Venn diagram shows the overlap between the proteins identified using different trypsin digestion durations. For each timepoint, the protein set is defined as the intersection of proteins identified in two biological replicates.

We further demonstrate here that this novel approach can shed light on the functions of under-studied ECM proteins by using as a case study the novel ECM protein SNED1 (Sushi, Nidogen and EGF Like Domains 1). We previously identified SNED1 in a proteomic screen comparing the ECM of poorly and highly metastatic mammary tumors [39]. Functionally we showed that SNED1 acts as a metastasis promoter, since knocking down SNED1’s expression significantly decreased metastatic dissemination. We also showed that the organization of fibrillar collagens within the tumor ECM was altered upon SNED1 knockdown [39], which led us to postulate that SNED1 may modulate ECM architecture. However, the precise mechanisms by which it could do so remain unknown. Using immunofluorescence microscopy, we have recently reported that SNED1 forms fibrils within the ECM [40]. Using, molecular modeling, we have predicted the interactome of SNED1 and found that SNED1 could potentially interact with 40 ECM components, including fibronectin, nidogens 1 and 2, and this may serve as the basis for SNED1’s incorporation in the ECM [40]. However, none of these interactions have been validated experimentally yet, and we still do not know the mechanisms underlying SNED1’s assembly in the ECM. Here, by applying time-lapsed proteomics to profile ECM produced *in-vitro* by cells overexpressing SNED1, we show that SNED1’s expression and presence in the ECM modulates ECM protein composition. Interestingly, we also show that SNED1 increases the accessibility of some ECM proteins to trypsin resulting in an increase sequence coverage for these proteins (*e*.*g*., thrombospondins 1 and 2, Efemp1, Efemp2, fibrillin 2) but also decreased accessibility of other ECM proteins to trypsin, resulting in a decrease sequence coverage for these proteins (*e*.*g*., nidogen 2, Tinagl1, Tgm2). While the mechanisms by which SNED1 modulates ECM composition and ECM protein folding remains to be discovered, this proof-of-concept study highlights the ability of time-lapsed proteomics to shed light on novel or unsuspected roles for ECM proteins.

## RESULTS

### Time-limited proteolysis increases protein identification and coverage

To test the impact of modifying trypsin digestion duration on protein identification, we chose to work with ECMs produced by mouse embryonic fibroblasts *in vitro* (Figure 1A). The generation of cell-derived matrices (CDMs) after decellularization of fibroblast cultures is highly reproducible [41] and thus offers a robust system to test and optimize ECM proteomic workflows. Once CDMs are obtained, they are partially solubilized and pre-digested with Lys-C (Figure 1A). CDMs are then either digested into peptides using a standard 18-hour tryptic digestion protocol [17,38] or using time-limited digestion (30 minutes, 2 hours, or 4 hours; Figure 1B). Peptides obtained were then characterized using LC-MS/MS (Supplementary Table 1, Supplementary Table 2). Overall, we observed that, time-limited digestions resulted in the highest absolute number of overall and matrisome spectra, while the proportion of matrisome spectra remained equivalent (Supplementary Tables 1 and 2).

In terms of protein identification, we detected, 77, 79, 88, and 76 ECM proteins across two biological replicates using 30-min, 2-hour, 4-hour, and 18-hour digestions, respectively (Figures 1C-D, Supplementary Tables 1 and 2). Across all digestions, about 2/3 of the proteins belong to the core matrisome (ECM glycoproteins, collagens, proteoglycans) while 1/3 are matrisome-associated proteins (ECM-affiliated proteins, ECM remodeling enzymes, or secreted factors). Comparison of the proteins identified at the different timepoints reveals that 68 proteins were identified across all timepoints (Figure 1D, Supplementary Table 3). In addition, 2 proteins (Col4a3 and Fgf2) were uniquely identified by the 30-minute digestion, while 9 proteins were uniquely identified by the longer 2-hour and 4-hour digestions (Mfge8, Vwa1, Thbs2, Lum, Anxa4, Anxa5, Loxl3, Egfl7 and S100a13), and an additional 6 only using the 4-hour digestion protocol. While the standard 18h digestion protocol uniquely identified 2 proteins (annexin 5 and S100a13), it resulted in an overall lower number of proteins identified (76 vs. 87 with the 4-hour digestion).

Beyond protein identification, we next sought to evaluate the impact limited proteolysis on sequence coverage. We found that limited digestions resulted in an increase in the percentage of sequence coverage for all categories of matrisome proteins as compared with a standard 18hour digestion (Table 1, Supplementary Table 2).

**Table 1.** Average sequence coverage achieved using different digestion durations for the different matrisome protein categories.

|  | Average % sequence coverage |  |  |  |
| --- | --- | --- | --- | --- |
|  | 30 min | 2h | 4h | 18h |
| **ECM Glycoproteins** | 21.5% | 25.2% | 22.9% | 15.9% |
| **Collagens** | 9.7% | 8.2% | 10.0% | 9.0% |
| **Proteoglycans** | 34.8% | 35.1% | 37.0% | 26.5% |
| **ECM-affiliated** | 14.7% | 14.7% | 12.8% | 8.0% |
| **ECM Regulators** | 18.0% | 19.9% | 19.4% | 10.3% |
| **Secreted Factors** | 12.7% | 11.2% | 9.0% | 11.2% |

### Sequential tryptic digestion as a mean to fractionate peptides and further improve protein identification and sequence coverage

We next sought to perform a time-lapsed digestion where peptides are collected sequentially, at different timepoints, over an 18-hour period (Figure 2A). To compare the two digestion protocols (single timepoints vs time-lapsed), we first evaluated their output in terms of protein identification. Using the time-lapse digestion protocol, we identified 76, 75, 67, and 41 ECM proteins in two biological replicates, at the 30-min, 2-hour, 4-hour, and 18-hour timepoints, respectively (Figure 2B-C). We also observed that the absolute number of spectra and peptides decreased at later timepoints of the time-lapse digestion (Supplementary Table 1). This is expected, since easy-to-digest peptides are depleted from the samples at the earlier timepoints of the sequential digestion.

**Figure 2.**
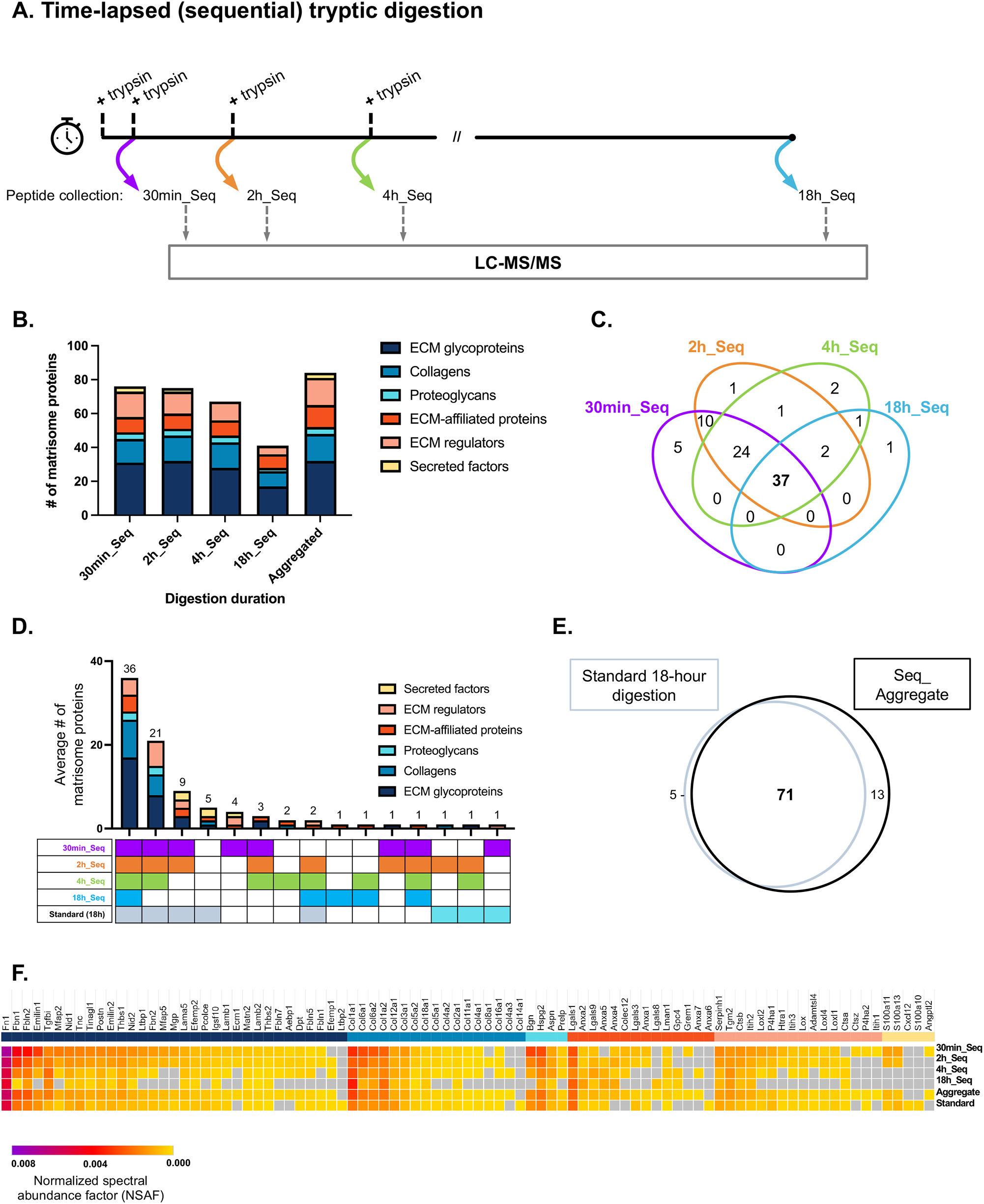
Identification of ECM proteins using time-lapsed trypsin digestion. **A**. Schematic representation of the sequential trypsin digestion protocol to achieve time-lapsed proteolysis. **B**. Bar graph depicts, for each matrisome category, the number of ECM proteins identified in two biological replicates using a time-lapsed digestion protocol. Aggregation of the data from all 4 timepoints of the time-lapsed digestion is also provided. For this, the average number of proteins identified across 2 biological replicates, is obtained first by defining proteins identified at any of the four digestion timepoints for each replicate, and then averaging these numbers (*see also Supplementary Tables 2 and 3*). **C**. Venn diagram shows the overlap between the proteins identified at each timepoint of the time-lapsed digestion protocol. For each timepoint, the protein set is defined as the intersection of proteins identified in two biological replicates. **D**. Matrisome proteins identified at each of the four sequential timepoints or the standard 18-hour single timepoint digestions are compared, and the intersections are visualized using an UpSet plot (*see also Supplementary Tables 2 and 3*). **E**. Venn Diagram illustrates the comparison between matrisome proteins identified using the standard 18-hour digestion protocol and upon aggregation of the data from the 4 sequential digestion timepoints, both lists being defined as the ensemble of matrisome proteins identified in two biological replicates. **F**. Heatmap illustrates the normalized spectral abundance factor (NSAF) obtained for each matrisome protein using the time-lapse trypsin digestion protocol or the standard single timepoint (18-hour) digestion protocol.

Aggregation of the data from the sequential digestions identified an average of 87 ECM proteins (Figure 2B, *Aggregated*), as compared to the 76 proteins identified using the standard 18-hour digestion. The comparison of the proteins identified in two biological replicates showed a smaller overlap of 37 proteins across all timepoints (Figure 2C) compared to that of the single timepoint digestions (Figure 1D). This overlap consists of 17 ECM glycoproteins, nine collagens, two proteoglycans, five ECM-affiliated proteins, and 4 ECM regulators (Figure 2C, Supplementary Table 3). 24 matrisome proteins were identified in samples from the first three timepoints, ten matrisome proteins in samples from the first two timepoints. We also found that the first timepoint of the time-lapse digestion (30 minutes) contributed the most to the aggregate list of ECM proteins identified as an additional five matrisome proteins (Grem1, Ctsz, P4ha2, Itih1, and Angptl2) were uniquely identified at this timepoint (Figure 2C, Figure 2F).

Of note, proteins that tended to vary more between protocols, timepoints, and replicates belonged to the matrisome-associated classification [17]. These proteins are expected to be present in lower abundance in CDMs than the more structural core matrisome proteins. For this reason, a short digestion cannot singlehandedly capture the complexity of CDM composition as it misses proteins present in lower abundance. This is further demonstrated by the five-way comparison of datasets obtained with the four timepoints of the time-lapsed digestion or the standard 18-hour digestion (Figure 2D, Supplementary Table 3). The comparison yielded 36 matrisome proteins identified in all samples. These consisted of 17 ECM glycoproteins, 9 collagens, 2 proteoglycans, 4 ECM-affiliated proteins, and 4 ECM regulators. It also showed how later timepoints of the time-lapse digestion are contributing complementary datapoints to the 30-minute digestion (Figure 2E, Supplementary Table 3). The other timepoints captured four matrisome proteins identified using the standard digestion protocol that otherwise would have been missed, and further add the identification of four matrisome proteins that the standard digestion did not identify. We therefore defined the aggregate of identified proteins from the sequential digestion as the union of four timepoints, which creates a list greater than that of the classical digestion, with 13 uniquely identified proteins (Figure 2E).

To further refine the characterization of the matrisome produced by MEFs *in vitro*, we compared the relative abundances of the proteins identified using normalized spectral abundance factor (NSAF), a semi-quantitative method based on spectral counts normalized by protein size [42,43]. The most abundant matrisome proteins found in MEF CDMs are fibronectin (Fn1), fibrillin 1 (Fbn1), fibulin 2 (Fbln2) and the elastin microfibril interface-located protein 1 (Emilin 1) for the glycoproteins and type I and type VI collagens (Figure 2F, Supplementary Table 7). Interestingly, we also detected basement membrane proteins including nidogens 1 and 2 (Nid1 and Nid2), perlecan (Hspg2) and type IV collagens.

As anticipated, we observed that the abundances of most proteins decreased with time during the time-lapsed digestion (Figure 2B, Supplementary Table 1, Supplementary Table 2). We also observed that the relative abundance of some proteins increased with longer digestions (Tgfbi, Tgm2, Col1a1, Col1a2, Col18a1, Lgals1, Anxa5, Anxa4). This may suggest that these proteins require more extensive denaturation and digestion to release peptides.

### Time-lapsed digestion increases matrisome protein sequence coverage

After evaluating the impact of the two digestion protocols on protein identification, we sought to compare the methods in terms of protein sequence coverage, in other word, the percentage of a given protein sequence matched (or “covered”) by peptides detected by mass spectrometry. We found that proteins showed higher coverage at earlier digestion timepoints (Supplementary Figure 1A). In particular, average coverage of proteins with peptides detected at the 2-hour timepoint of the sequential digestion was the highest of the four timepoints and surpassed that of the standard 18-hour digestion (Supplementary Figure 1B). However, by combining the identified peptide sequences of earlier timepoints with those of the next timepoint (aggregation), we observed that the coverage was significantly higher over time (Figures 3A, 3B, Supplementary Figure 1B). This increase in coverage over time for most proteins suggests that some of the peptide sequences identified are unique to the later timepoints of the time-lapsed digestion, thus increasing the cumulative coverage with time (Figure 3A).

**Figure 3.**
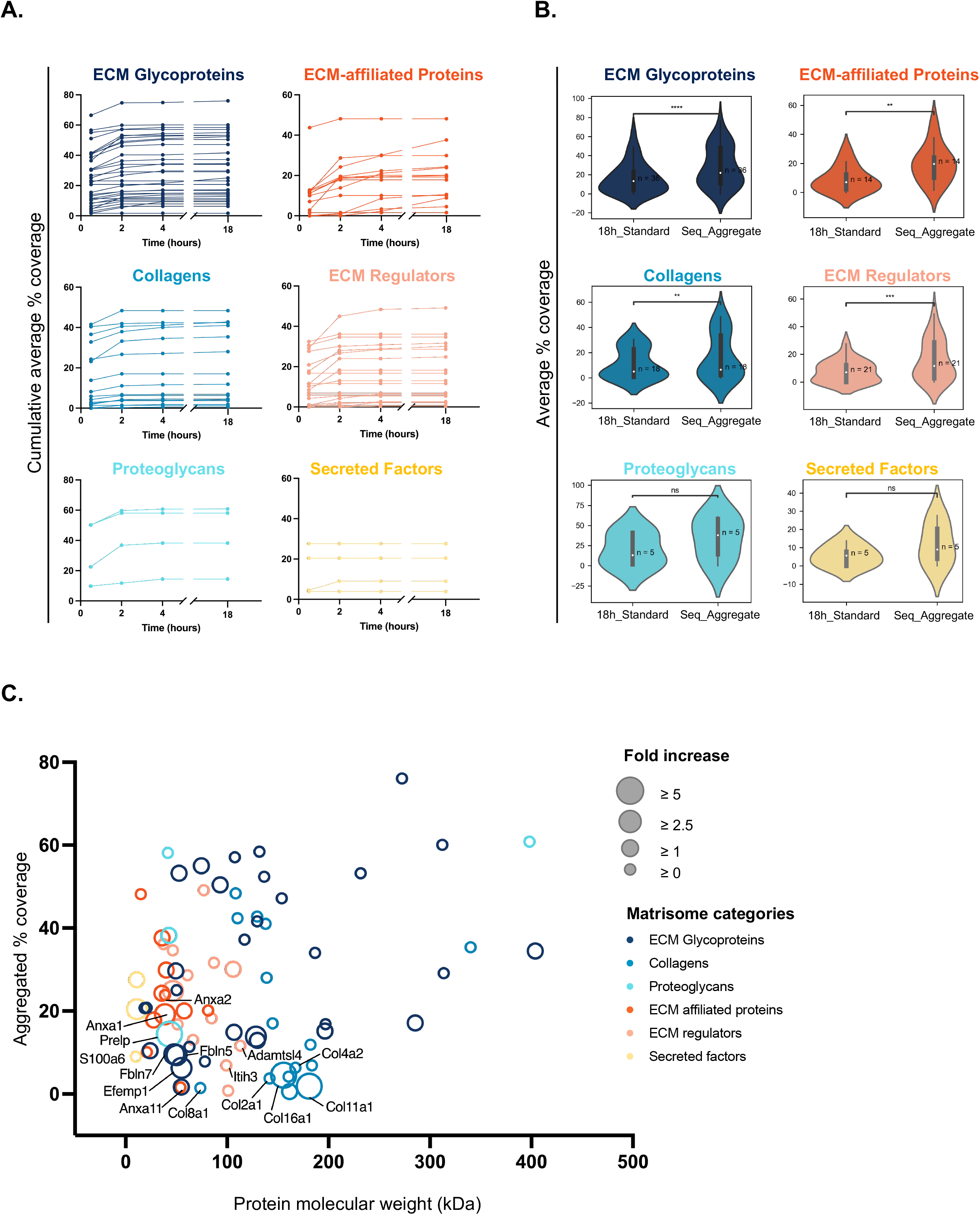
Aggregation of data from sequential tryptic digestion timepoints results in enhanced sequence coverage. **A**. Each line illustrates the cumulative average sequence coverage percentage at each timepoint for a given matrisome protein (*see also Supplementary Table 2*). **B**. Violin plots compare, for each matrisome category, the distribution of average protein sequence coverage obtained with a standard 18-hour digestion protocol or by aggregating data from the 4 timepoints of the time-lapsed digestion. Median and interquartile range are indicated by the white dots and grey bars, respectively. Wilcoxon signed-rank test was used to compare the distribution, and p-value was reported after Bonferroni adjustments were made (*p<0.05, **p<0.01, ***p<0.001, ****p<0.0001, ns: not significant). **C**. Scatter plot represents the average aggregated percentage of sequence coverage for each protein detected (y-axis) as a function of protein molecular weight (kDa, x-axis). The diameter of each dot represents the fold-change in coverage as compared to the standard 18-hour single timepoint digestion method, and the proteins with changes above 90^th^ and below 10^th^ percentile are annotated (*see Table 2 and Supplementary Table 2*).

To evaluate whether a particular matrisome category of proteins benefitted more from this approach, we compared the protein coverages obtained using the standard 18-hour digestion or after aggregating coverage data of the time-lapsed protocol. We observed a shift in the distribution of coverages particularly for ECM glycoproteins (p < 0.001), ECM-affiliated proteins (p < 0.01), and ECM Regulators (p < 0.01) (Figure 3B). The increase in cumulative coverage is most noticeable with the ECM-affiliated proteins and ECM regulators, which are less abundant, and therefore would have benefitted the most according to our hypothesis (Figure 3A, 3B).

To rule out that the increase in the sequence coverage seen upon data aggregation was due to the increase in the number of datasets collated, we aggregated four datasets obtained using the standard 18-hour digestion method (*i*.*e*., four files) and compared the average aggregated coverage to the average aggregated coverage obtained by integrating data from the four distinct digestion timepoints (Supplementary Figure 1C). This confirmed that the increased percentage of sequence coverage is indeed due to the benefit of the time-lapsed digestion method and the release of different peptides over time, and not the collation of datasets.

To evaluate the contribution and potential benefit of each timepoint to the increased coverage attained with the time-lapsed digestion protocol, we calculated the coverage with increasing extent of aggregation—30-min to 2-hour, 30-min to 4-hour, 30-min to 18-hour—and evaluated their correlations to one another (Supplementary Figure 2). While the differences between the cumulative aggregated coverage of 4-hour and that of the 18-hour were minimal, as denoted by Pearson’s correlation coefficient (r > 0.97), the greatest differences were seen between the 30-min unaggregated coverages and the cumulative aggregated coverage of 18-hour (r = 0.93) (Supplementary Figure 2). Upon calculation of cumulative coverage, we found that the highest coverage was achieved for fibronectin (76 %, while fibronectin’s coverage was of ∼60% using an 18-h standard digestion protocol, *see* Table 2 and Supplementary Table 2), followed by perlecan (60 %, while fibronectin’s coverage was of ∼46% using an 18-h standard digestion protocol, *see* Table 1 and Supplementary Table 2).

**Table 2.**
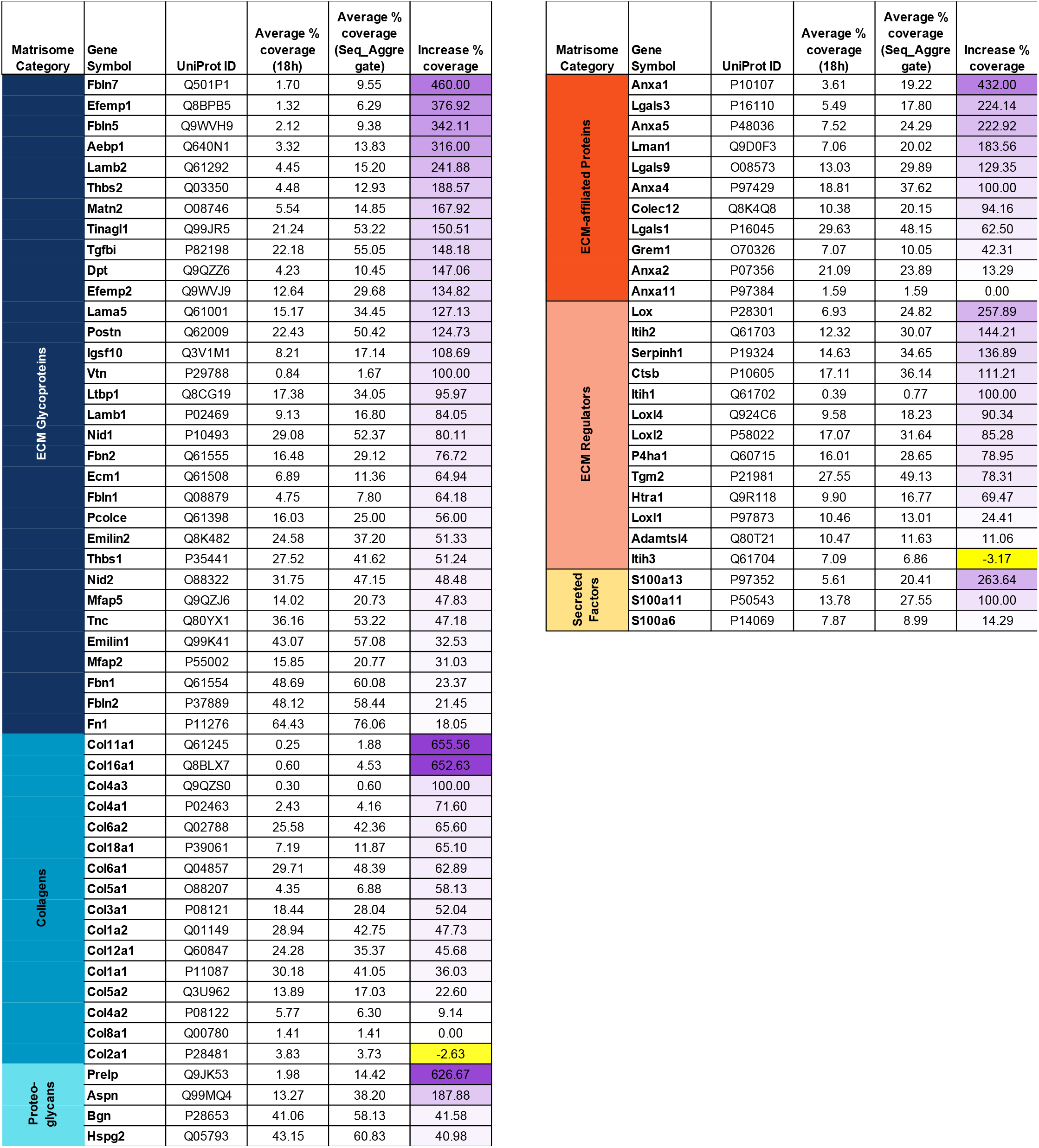
Average sequence coverage achieved using an 18-hour or a time-lapsed tryptic digestion

To further identify the proteins that benefitted the most from the increase in coverage achieved by employing a time-lapsed digestion protocol, we calculated the ratio of aggregated coverage to standard digestion coverage, for each protein (Table 2). We found that type XI and type XVI collagens (Col11a1 and Col16a1), the proteoglycan prolargin (Prelp), and several fibulins (Fbln7, Efemp1, Fbln5) saw the largest increase in coverage. These changes in coverage were further plotted with respect to protein molecular weight, to evaluate whether a protein’s size affected the observed coverage increase (Figure 3C, Supplementary Table 2). While a greater number of ECM glycoproteins saw an improvement in their coverage (∼50 % of all identified ECM glycoproteins), the extent of improvement was similar to those of the proteoglycans, ECM-affiliated proteins, and ECM regulators (Figure 3C, Table 2). The overlap of the distribution with respect to size and coverage observed between the most and least affected proteins suggested no particular effects of these parameters on the increased coverage upon data aggregation (Figure 3C).

In conclusion, we demonstrate that, while performing time-lapsed tryptic digestion and aggregating data results in an only moderate increase in protein identification, it permits a more accurate estimation of relative protein abundance and leads to a significant increase in protein sequence coverage. Because the time-lapsed digestion protocol results in the release of peptides over time, we can also obtain information on protein digestibility and accessibility and then infer information about protein folding. The first draft of the MEF matrisome protein accessibility map provided here, thus represent a first step towards obtaining a detailed topographical map of ECM.

### Identification of SNED1-dependent changes in ECM protein composition and abundance

Our next step was to test whether this new protocol could help us gain new knowledge on the ECM. We previously reported changes in the organization of the ECM of mammary tumors in presence of SNED1 which correlated with tumor’s metastatic potential [39]. More recently, we have shown that SNED1 is incorporated in fibrillar structures within the ECM meshwork (Figure 4A, [40]), yet, we do not know how SNED1 assembled into an ECM and how it modulates ECM architecture. We thus sought to use the newly developed time-lapsed proteomic workflow presented here to gain insight into the possible roles of this protein on ECM composition and ECM protein abundance and folding. To do so, we analyzed CDMs produced by *Sned1*^*KO*^ MEFs over-expressing GFP-tagged SNED1 and compared it to the CDMs produced *Sned1*^*KO*^ MEFs over-expressing GFP alone (control; *see above*). Quality-control analysis of the dataset obtained, on the model of what we presented above, revealed a similar pattern in terms of protein identification using single timepoint digestions or time-lapsed digestion (Supplementary Figure 3, Supplementary Tables 4 and 5).

**Figure 4.**
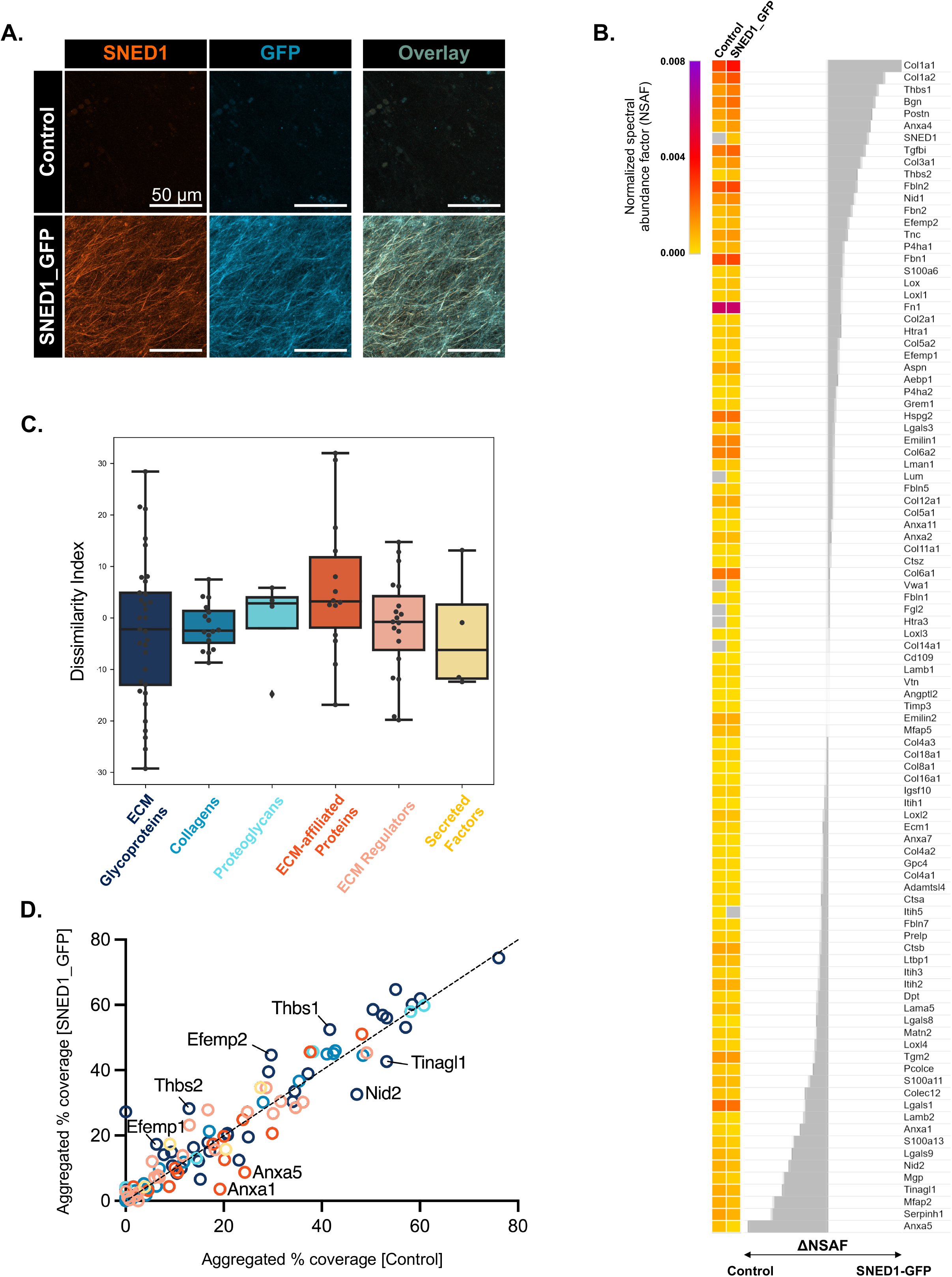
The ECM protein SNED1 modulates MEF matrisome composition and coverage. **A**. Immunofluorescence images of decellularized CDMs produced by immortalized mouse embryonic fibroblasts overexpressing GFP alone (control, top panels) or SNED1_GFP (bottom panels) co-stained with anti-SNED1 (orange, left panels) and anti-GFP antibodies (light blue, middle panels), producing an overlay (right panels). Scale bars: 50 μm. **B**. Heatmap represents the aggregated normalized spectral abundance factor (NSAF) of proteins identified in CDMs produced by control or *SNED1_GFP*-overexpressing MEFs. Difference in abundance is calculated by subtracting the relative abundance of proteins of control CDMs from those of SNED1-containing CDMs (*see Supplementary Table 4)*. Proteins are sorted in descending order of differential abundance. **C**. A custom dissimilarity index was calculated to represent changes in protein sequence coverage in the aggregated dataset of the 4 timepoints of the time-lapsed digestion of control and SNED1-containing CDMs. Proteins depicted by dots toward the top of the boxes show matrisome proteins whose sequence coverage is lower when SNED1 is present, while proteins depicted by dots toward the bottom show the opposite. **D**. Scatter plot represents the relation between the average sequence coverage of all matrisome proteins detected in both control and SNED1-containing CDMs. Proteins found above the diagonal (*e*.*g*., Thbs1, Thbs2, Efemp2) are proteins whose coverage increase in SNED1-containing CDMs, while proteins found below the diagonal (*e*.*g*., Nid2, Tinagl1) are proteins whose sequence coverage is decreased in SNED1-containing CDMs (*see Table 3 and Supplementary Table 4*). Proteins with dissimilarity index above 95^th^ and below 5^th^ percentile are annotated.

As described above, we could define the composition of CDMs containing or not SNED1 as the list of proteins at the intersection of two biological replicates for each condition (Supplementary Table 6) and identified proteins whose detection changed as a function of SNED1’s expression. Doing so revealed that the *α*3 chain of type IV collagen (Col4a3), the *α*1 chain of type VIII collagen (Col8a1), and Itih1 were not detected in the control CDMs, while S100a6, was only identified in SNED1-containing CDMs (Supplementary Table 6).

We next sought to evaluate whether the presence of SNED1 altered the abundance of certain ECM proteins. To do so, we calculated the NSAF for all proteins detected in SNED1-containing CDMs (Supplementary Figure 3G, Supplementary Table 7) and compared these values with those obtained for control CDMs (Figure 4B, Supplementary Table 7). We found that type I collagen (chains *α*1 and *α*2), the glycoproteins thrombospondin 1 and periostin (Postn), the proteoglycan biglycan (Bgn), and annexin A4 were more abundant in SNED1-containing CDMs. In contrast, the glycoproteins Mfap2, Tinagl1, Mgp, and Nid2 and the Anxa5, Serpinh1, were present in lower abundance, in SNED1-containing CDMs as compared to control.

### Changes in ECM protein profiles upon expression of SNED1

To gain insight into the possible role of SNED1 in modulating ECM architecture, we exploited the coverage data obtained by proteomics. Indeed, protein digestibility by trypsin is directly linked to how accessible lysine and arginine residues are, and this is linked to protein folding and protein-protein interactions, two parameters that can alter exposure of these residues. To capture this, we defined a dissimilarity index reflecting the magnitude of the differences in coverage (Figure 4C, Table 3, *see Materials and Methods*). Using this index, we identified proteins with increased coverage in CDMs containing SNED1, including thrombospondins 1 and 2 (Thbs1 and Thbs2) and EGF containing fibulin extracellular matrix protein 2 (Efemp2, also called fibulin 4), while proteins with decreased coverage in SNED1-containing CDMs included of emilin1, Tinagl1, nidogen 2, galectin 9, and annexin A5 (Figure 4D).

**Table 3.**
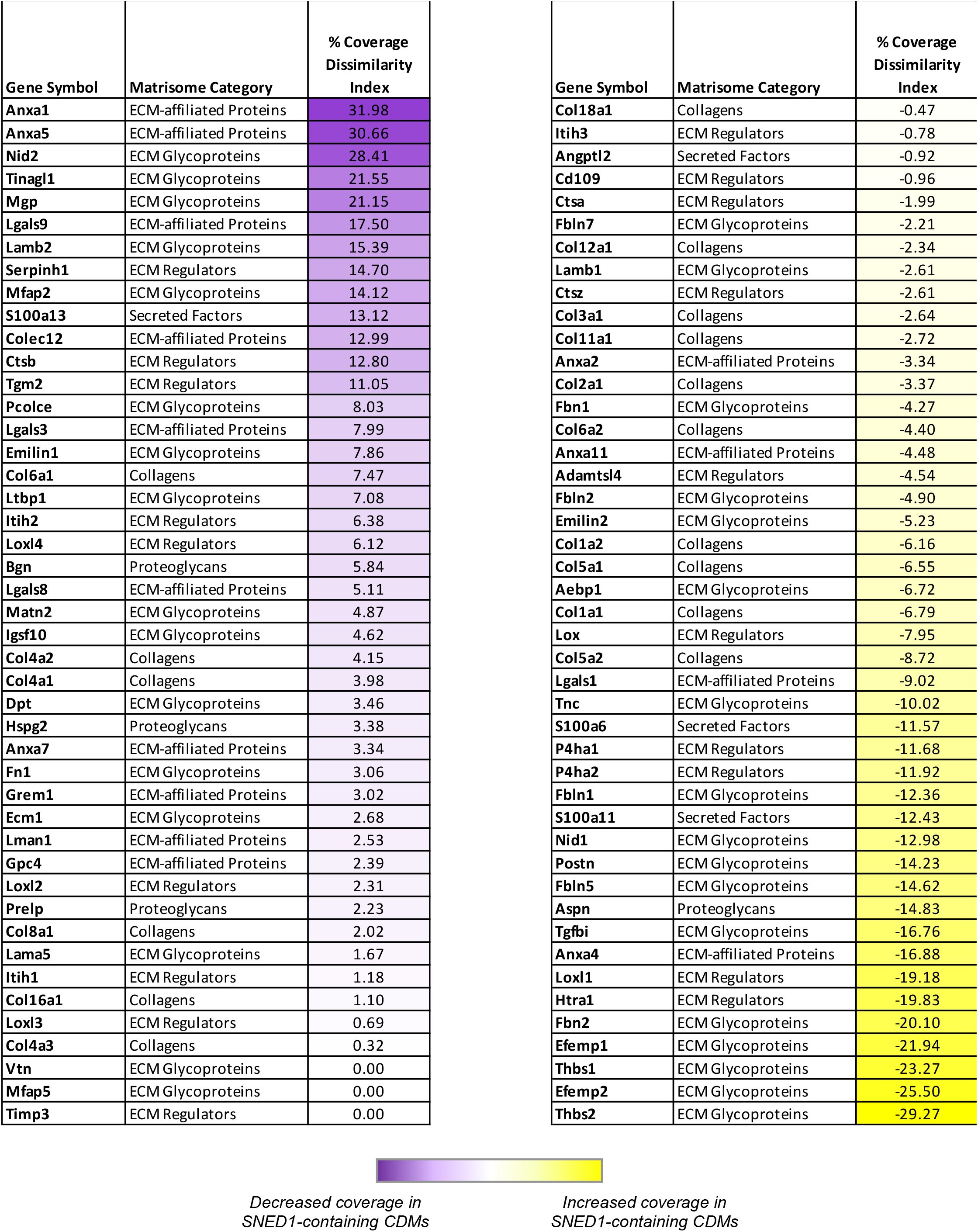
Sequence coverage dissimilarity index

Since an increase in protein abundance could result in an artificial increase in coverage, we next sought to identify proteins detected with increased coverage without differential abundance. With this approach, proteins such as Fbln1, Col4a1 can be considered as those whose increase or decrease in coverage, respectively, is independent of their relative abundance. On the other hand, proteins such as Col1a1, Col1a2, Fbln2, Fbn1, Col2a1 are identified with increased abundance without increase in coverage.

Altogether, our results support the conclusion that SNED1’s expression modulates the composition and coverage of the ECM produced by fibroblasts. It will be interesting in the future to identify the mechanisms leading to these changes (*see Discussion*).

### Sequential release of peptides provides insight on protein accessibility to trypsin

Perhaps the most significant innovation brought by the time-lapsed digestion protocol presented here is its spatio-temporal resolution. As discussed above, protein digestibility reflects the surface exposure of a protein, and is directly linked to protein folding and protein-protein interactions. Thus time-lapsed peptide mapping can identify protein regions readily exposed and accessible to trypsin (peptides matching these regions would be released at early digestion timepoints) and regions of proteins hard to access, either because of extensive post-translational modifications, such as O-glycosylations hindering accessibility, or because the engagement of protein-protein interactions that are not disrupted by the partial denaturation to which samples are submitted.

#### Digestion kinetics and topographical information of SNED1

As of today, we do not have any experimentally-validated information on the folding of SNED1 when incorporated in the ECM. To obtain topographical information on SNED1, we assessed the diversity of peptides generated over the course of the time-lapsed digestion. Overall, we found that the aggregated SNED1 coverage yielded 27.32 %, a significant increase from the 13.8 % coverage obtained using the 18h standard digestion protocol (Supplementary Table 4). It is also a substantial increase from the ∼5% coverage obtained by aggregating data from MatrisomeDB, a database reporting proteomics data on insoluble ECMs from tissues [35], and an increase from the 24.13% coverage reported in ProteomicsDB, with the caveat that this database compiles global proteomic datasets and that for the most part includes proteomic analysis of soluble protein samples [44]. We observed that the number of peptide spectrum matches (PSMs) was the highest for the 30-minute timepoint of the time-lapse trypsin digestion. The most peptides at this timepoint were in the CCP/Sushi domain (SMART SM000032) and in the first FN3 (SMART SM000060) domain, followed by peptides in the NIDO domain (SMART SM000539) (Figure 5A, Supplementary Table 8). Over time, the number of PSMs decreased, yet we found that each timepoint contributed unique peptides. In particular, we saw that a large number of peptides mapped to the second FN3 domain at the later digestion timepoints, with six and five PSMs for the 4-hour and 18-hour timepoints, respectively, which suggests that this domain gets exposed over time as the tryptic digestion progresses.

**Figure 5.**
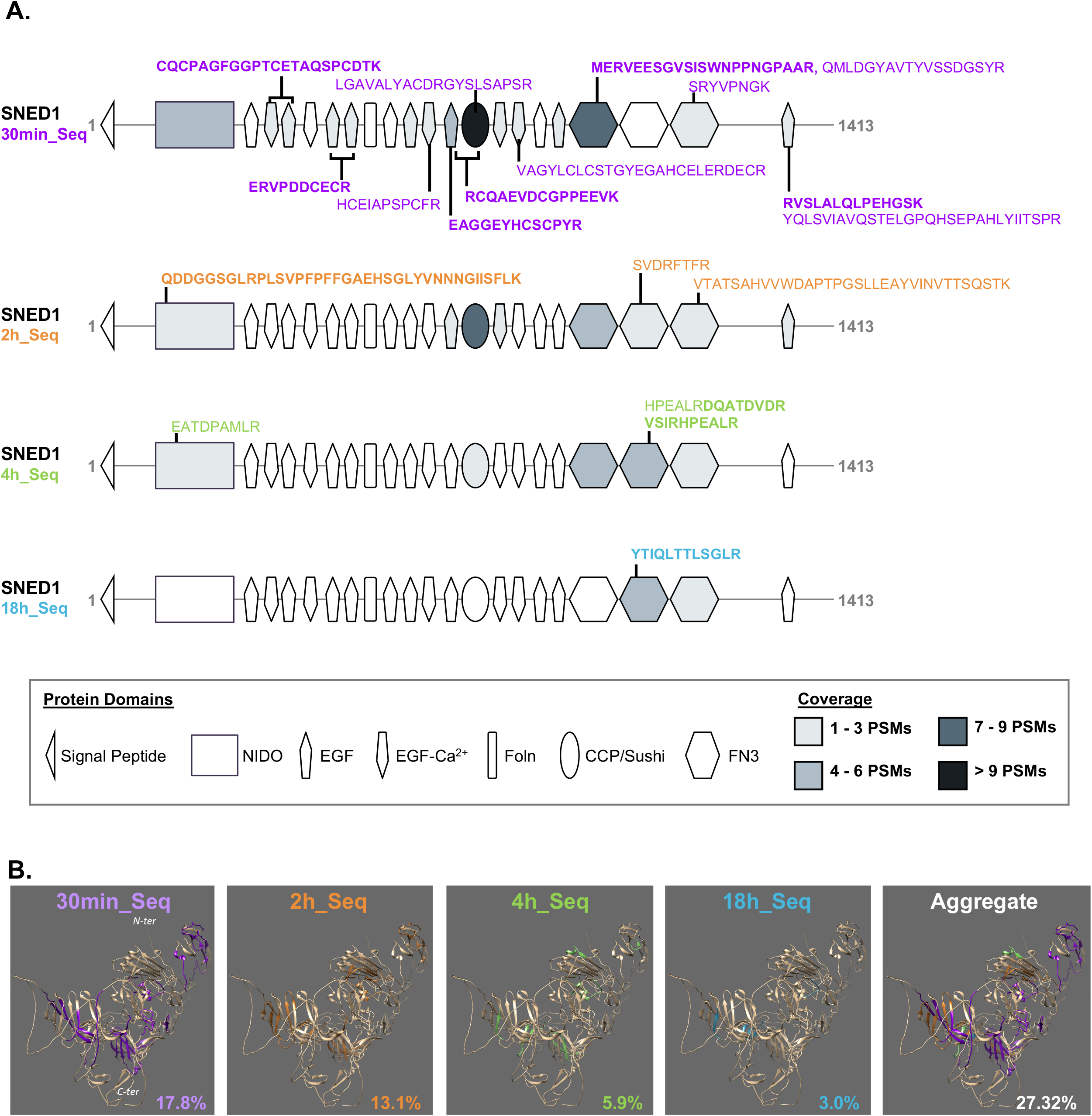
Time-lapsed peptide mapping of SNED1. **A**. Mapping of peptides detected at each of the timepoints of the time-lapsed digestion on schematic representation of the domain-based organization of SNED1 (UniProt Q8TER0). Each peptide is mapped using the start and end position of each peptide, based on information provided by the Simple Modular Architecture Research Tool (SMART, http://smart.embl-heidelberg.de/). The color scale from light to dark grey represents, for each domain, the number of peptide-spectrum matches (PSMs). Peptide sequences depicted are the ones uniquely identified at a given timepoint. Bolded sequences correspond to peptides detected in two biological replicates. **B**. 3-D mapping of detected peptides on the AlphaFold-predicted model of SNED1 (https://alphafold.ebi.ac.uk/entry/Q8TER0). Peptides are color-coded to indicate at which digestion timepoints they were detected. For the aggregate illustration the timepoint was assigned to the timepoint of first appearance.

While the structure of SNED1 has not been resolved yet, we can turn to molecular modeling to gain insight into the 3-dimensional folding of the protein. Here, we retrieved the 3-D structure of SNED1 predicted by AlphaFold [45] (https://alphafold.ebi.ac.uk/entry/Q8TER0) and mapped, in a color-coded manner, the peptides identified at each timepoint of the time-lapsed digestion (Figure 5B, Supplementary Table 8, Supplementary File 2). While the folding of some regions of the protein is only predicted with low confidence and while it is likely that SNED1’s folding is different when it is assembled in the ECM, this is a first attempt at overlaying experimental data obtained by time-lapsed proteomics on a theoretical model of an ECM protein.

#### Peptide mapping of nidogen 2 whose coverage decreased upon SNED1presence

To illustrate how SNED1 presence in the ECM influenced the coverage of ECM proteins, we performed a similar peptide mapping for nidogen 2. We found that Nidogen 2 (Nid2) had a higher sequence coverage in the CDMs lacking SNED1 (Figure 4C, Figure 6A, Table 3). The G2F domain was identified with the highest number of peptides, with 45 in the SNED1-lacking CDM and 36 in the SNED1-containing CDM. This was followed by the NIDO domain which was identified with 33 spectra in the SNED1-lacking CDM and 21 spectra in the SNED1-containing CDM. The 3-D model of Nid2 predicted by AlphaFold was further used to overlay peptide identified at each timepoint of the time-lapsed digestion. This revealed the precise location of peptides detected differentially in presence or absence of SNED1 (Figure 6B, Supplementary Files 3 and 4). We can speculate that peptides released at later timepoints of the time-lapses digestion are part of protein domains not as readily accessible to tryptic digestion, and possibly because these domains are involved in protein-protein interactions, protecting them from tryptic digestion. Interestingly, nidogen 2 was one of the proteins we predicted to interact with SNED1 by molecular modeling [40], supporting the possibility that protein-protein interaction between SNED1 and nidogen 2 is leading to the decreased coverage of nidogen 2 in presence of SNED1. The fact that the presence of SNED1 modulates trypsin accessibility of several ECM proteins, suggests that this novel protein of the ECM plays a key role in modulating the assembly and organization of the ECM.

**Figure 6.**
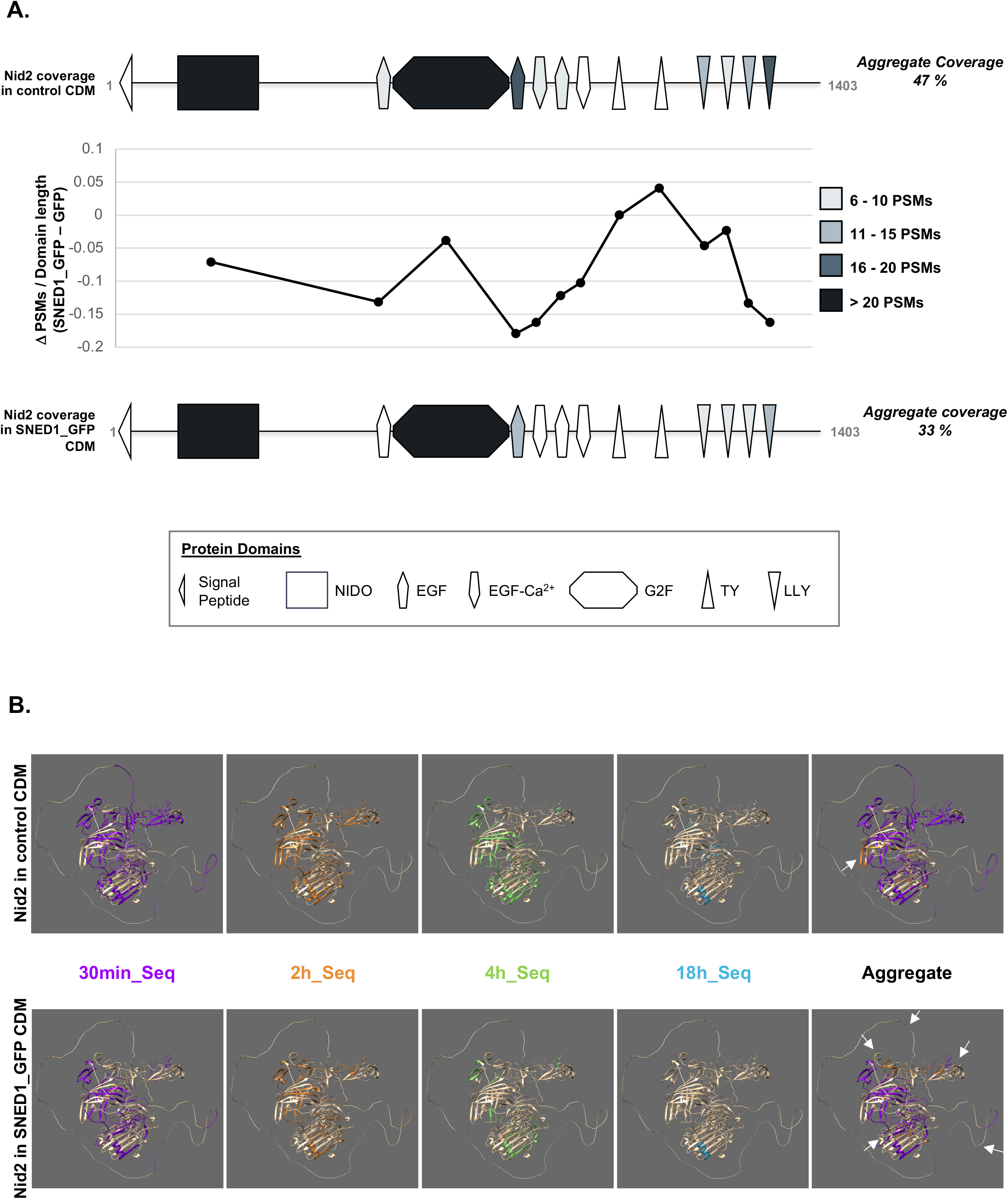
Nidogen-2 sequence coverage decreases upon SNED1 overexpression. **A**. Representation of peptide mapping of nidogen-2 (Nid2, UniProt O88322). Upper panel represents the mapping of Nid2 peptides detected in control CDM samples; lower panel represents the mapping of Nid2 peptides detected in SNED1-containing CDM samples. The color scale from light to dark grey represents, for each domain, the number of peptide-spectrum matches (PSMs). The line plot illustrates the difference in numbers of PSM detected in control or SNED1-containing CDMs and normalized by domain length. **B**. 3-D mapping of detected peptides on the AlphaFold-predicted model of nidogen-2 (Nid2; https://alphafold.ebi.ac.uk/entry/O88322). Upper panel represents the 3-D mapping of Nid2 peptides detected in control CDMs; lower panel represents the 3-D mapping of Nid2 peptides detected in SNED1-containing CDMs. Each peptide color-coded to indicate the timepoint at which it was detected. For the aggregate illustration, the timepoint was assigned to the timepoint of first appearance.

#### Peptide mapping of thrombospondin 1 whose coverage increased upon SNED1presence

In contrast with what we observed for nidogen 2, thrombospondin 1 saw its sequence coverage increased when SNED1 is present in the ECM (Figure 4C, Table 3, Figure 7A). We found that the thrombospondin N-terminal like domain contained the highest number of PSMs, 102 in SNED1-containing CDMs, compared to 78 counts in SNED1-lacking CDMs. To better capture differences, on the model of what has been proposed by Eckersley and others [Eckersley et al 2020], PSM counts were normalized by the amino-acid length of each domain (Figure 7A).

**Figure 7.**
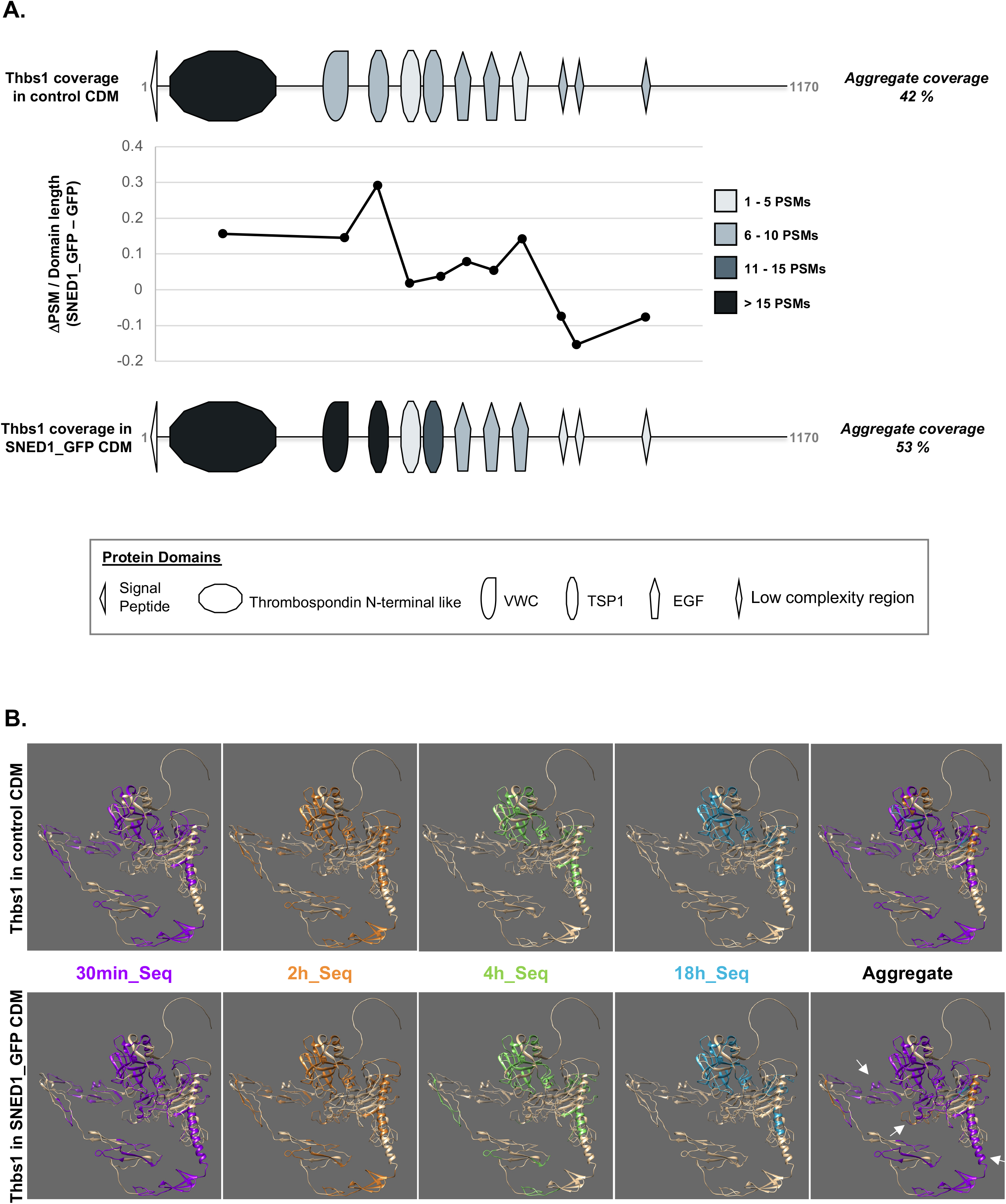
Thrombospondin-1 sequence coverage is increased upon SNED1 presence in the ECM. **A**. Representation of peptide mapping of thrombospondin-1 (Thbs1, UniProt P35441). Upper panel represents the mapping of Thbs1 peptides detected in control CDM samples; lower panel represents the mapping of Thbs1 peptides detected in SNED1-containing CDM samples. The color scale from light to dark grey represents, for each domain, the number of peptide-spectrum matches (PSMs). The line plot illustrates the difference in numbers of PSM detected in control or SNED1-containing CDMs normalized by domain length. **B**. 3-D mapping of detected peptides on the AlphaFold-predicted model of thrombospondin-1 (Thbs1; https://alphafold.ebi.ac.uk/entry/P35441). Upper panel represents the 3-D mapping of Thbs1 peptides detected in control CDMs; lower panel represents the 3-D mapping of Thbs1 peptides detected in SNED1-containing CDMs. Each peptide is color-coded to indicate the timepoint at which it was detected. For the aggregate illustration, the timepoint was assigned to the timepoint of first appearance.

The 3-D model of Thbs1 predicted by AlphaFold was further used to overlay peptide identified at each timepoint of the time-lapsed digestion (Figure 7B, Supplementary Files 5 and 6). The resulting annotation showed a large overlap of peptide sequences identified across the different timepoints, revealing that most of the peptides were obtained with a short 30-minute digestion. Further analysis of the distribution of the peptides identified in SNED1-lacking CDMs with those identified in the SNED1-containing CDMs, revealed their precise topography (Figure 7B, white arrows). One hypothesis to explain how the presence of SNED1 in the ECM results in an increased in the sequence coverage of a protein, like thrombospondin 1, is if it directly or indirectly induced a conformational change on this protein, either by binding to it, or by modulating the forces exerted on it, which would, in turn, increase the exposure of this protein to trypsin. Future studies will be aimed at exploring this hypothesis.

## DISCUSSION

In this study, we report the development of time-lapsed digestion coupled to mass spectrometry to enhance ECM protein identification and sequence coverage, and gain insight into ECM protein folding in the context of an assembled ECM meshwork. By comparing MS output obtained from time-limited tryptic digestions, we found that shorter digestions are sufficient to identify the same number, if not more, than the gold standard overnight trypsin digestion protocol. We further demonstrate the significant benefit of conducting time-lapsed digestion and aggregating output in terms of protein identification and coverage over a standard overnight trypsin digestion protocol. Last, we highlight how time-lapsed proteomic data can be leveraged via 1-D and 3-D peptide mapping to gain insights into ECM protein folding and identification of possible sites of protein-protein interactions.

Modulation of the duration of the trypsin digestion is only one of the parameters of a multi-step proteomic workflow that can be modified. While we demonstrated its benefits, in particular in terms of sequence coverage, there is still a dark side to the matrisome that has yet to be illuminated. We were the first to propose and show that hard-to-digest ECM proteins would benefit from sequential multi-proteases digestion (in our case LysC + trypsin) [17], this combination is now broadly adopted. On the model of what has been developed to achieve higher coverage of the intracellular proteome [46–49], and based on the findings reported here, we propose that a sequential (or simultaneous) digestion with *multiple* proteases, beyond the use of the LysC-trypsin pair, could achieve a deeper matrisome coverage. However, this poses challenges including the optimization of reaction conditions to ensure protease compatibility and downstream computational analysis to support multiplexed digestion for accurate peptide identification.

The complex structure of the ECM meshwork mediated by protein-protein interactions that are dependent on proper protein folding, is crucial to its function. Conventional structural proteomic approaches such as cross-linking mass spectrometry (XL-MS, [50]) are not suitable to study ECM as crosslinking reagents will further harden already hard-to-digest ECM proteins. On the other hand, classical limited-proteolysis-coupled mass spectrometry method previously developed is devised to handle soluble proteins [37]. While the approach developed here can shed light on proteins’ accessibility in the assembled state of the ECM, translated as the proteins’ digestibility by trypsin, it is worth noting that it is applied here to samples that have been partially denatured. Our next step will be to attempt to work on native ECM samples to conduct a true topographical survey of the ECM using proteomics. In addition, in native conditions, protein domains engaged in protein-protein interactions will be inaccessible and as a result resistant to proteolysis. Peptide mapping should thus reveal, more directly, a map of potential protein-protein interaction sites at the ECM scale.

As a proof-of-concept, we applied the newly devised time-lapsed digestion protocol to compare the ECM produced by *Sned1*^*KO*^ or SNED1-overexpressing mouse embryonic fibroblasts. The *Sned1* gene was clones nearly two decades ago [51], yet, until very recently, we did not know anything about the physiological or pathological roles of SNED1. We became interested in studying SNED1 after discovering its role as a promoter of mammary tumor metastasis [39]. More recently, we reported the generation of a *Sned1* knockout mouse model and its essential role during development, since *Sned1* knockout led to early neonatal lethality, likely due to craniofacial malformations and its involvement in neural crest cell fate [52]. Based on sequence analysis, we categorized SNED1 as part of the core matrisome [17,30], before we knew that it could incorporate in the ECM and constitute one of its structural components, something we recently demonstrated experimentally [40]. However, how SNED1 assembles in the ECM and how it governs different pathophysiological processes remain unknown. Early observations looking at the ECM of tumors expressing or not SNED1suggested that SNED1 may play a role in modulating the tumor ECM [39]. We thus sought to use the newly developed time-lapsed proteomic pipeline to test this. Here we showed that overexpression of SNED1 by mouse embryonic fibroblasts indeed modulated ECM composition, as observed when calculating the relative abundance ECM proteins in cell-derived matrices produced by *Sned1*^*KO*^ or SNED1-overexpressing cells, and ECM protein accessibility to trypsin. Interestingly, several proteins whose abundance and/or whose coverage changed as a function of SNED1 are also predicted to be possible SNED1 interactors. These include fibronectin, and nidogen 2. Future work will be aimed at understanding the mechanisms leading to these changes and determine if, and if so, how, they relate to the pathophysiological process in which we have implicated SNED1.

Our view of the ECM has significantly changed over the past decades, from an inert scaffold to a remarkably complex and dynamic signaling platform. We now know that the ECM is actively remodeled over time during the processes of development, tissue aging, wound healing, and disease progression [4,7,8,10,12]. Changes in cell-ECM interactions, microenvironmental cues (*e*.*g*., tissue oxygenation levels), and environmental factors (*e*.*g*., smoke, UV) induce biochemical (*e*.*g*., glycation [53]) or structural (*e*.*g*., degradation, cross-linking) post-translational modifications, leading to alteration of the structure of individual ECM proteins and that of the ECM meshwork. Bottom-up proteomic and peptide mapping have contributed to appreciate and evaluate the extent of these changes for example during aging and photoaging of the skin [19– 21,27,54], fibrosis [11,26,27], or cancer progression [55], but much more remains to be discovered. As demonstrated here, we propose that applying time-lapsed proteolysis coupled to mass spectrometry can contribute to unravel the compositional and structural complexity of the ECM, a necessary first step toward deciphering ECM-dependent pathophysiological processes and the development of ECM-targeting therapeutic strategies.

## EXPERIMENTAL PROCEDURES

### Cell culture and generation of cell-derived matrices (CDMs)

Mouse embryonic fibroblasts (MEFs) were previously isolated from embryos of *Sned1*^LacZ-Neo/LacZ-Neo^ (*Sned1*^*KO*^) mice [52]. MEFs were first immortalized and then transduced with retroviral constructs to achieve stable expression of either the SNED1_GFP fusion protein (SNED1_GFP) or GFP alone (control), as previously described [40]. For all experiments, cells were cultured in DMEM (Corning, #10-017-CV) supplemented with 10 % fetal bovine serum (Sigma), and 2 mM glutamine (Corning).

To obtain cell derived matrices (CDMs), cells were cultured as described by Franco-Barraza, et al. [41]. In brief, cells were plated in 6-well plates at 475,000 cells/well. Upon cell reaching confluency (within 24 to 36 hours post seeding), the culture medium was replaced with fresh medium supplemented with 0.05 μg/mL of ascorbic acid (Sigma). Half of the medium was then replaced with fresh medium containing 0.1 μg/mL of ascorbic acid, every other day, for 7 days following the first ascorbic acid treatment. After 7 days of ascorbic acid treatment, cells were washed twice with warm D-PBS containing calcium and magnesium (Corning), and the ECM produced by the cells was decellularized using warmed extraction buffer (0.5 % Triton X-100 and 20 mM NH_4_OH in PBS) for 6 minutes. Decellularization efficiency was assessed by microscopic observation. The buffer containing cellular lysate was then removed and replaced with cold PBS. The obtained CDMs were cut in half using a scalpel and transferred from the cell culture plate to a conical tube. The CDMs were washed in PBS under constant inversion at 4 °C for 4 hours. The PBS was replaced after a centrifugation at 4,000 rpm for 8 minutes at 4 °C, and the wash was repeated once for 4 hours and then for 12 hours. Last, CDMs were pelleted by centrifugation at 13,000 rpm for 2 minutes at 4 °C, and stored at -80 °C.

For immunofluorescence visualization, cells were cultured on a glass coverslip coated with crosslinked gelatin as described by Harris *et al*. [56]. After decellularization (*see above*), CDMs were fixed with 4 % paraformaldehyde, blocked with 1 % BSA in 0.05 % Triton X-100, and incubated first with a polyclonal rabbit anti-SNED1 antibody (used at a 1 μg/mL concentration, previously described [40]) and a monoclonal mouse anti-GFP antibody (Abcam #ab1218, used at a 1:500 dilution), and then with a goat anti-rabbit antibody conjugated to AlexaFluor568 (Thermo Fisher A11036, used at 1:1000) and a goat anti-mouse antibody conjugated to AlexaFluor647 (Thermo Fisher A21236, used at 1:1000) for visualization. Coverslips were mounted using Fluoromount-G (Southern Biotechnology) and imaged using Zeiss confocal microscope LSM880.

### Protein solubilization and digestion into peptides

Each CDM pellet was resuspended in 100 mM NH_4_HCO_3_ solution containing 8 M urea and 10 mM dithiothreitol (DTT; Thermo Fisher) and incubated at 37 °C for 2 hours under agitation, for partial solubilization and reduction. Reduced thiol groups were alkylated using 40 mM iodoacetate (IAA; Thermo Fisher) for 30 minutes at room temperature in the dark. Urea concentration was brought to 2 M with 100 mM NH_4_HCO_3_ and samples were then de-glycosylated with PNGase F (New England Biolab), under agitation at 37 °C for 2 hours, and pre-digested with Lys-C (Thermo Fisher), under agitation at 37 °C for 2 hours, as previously described [17,38].

#### Single timepoint tryptic digestion

For single timepoint digestion (n = 4 for 2-hour and 18-hour timepoints; n = 2 for 30 min and 4-hour timepoints), trypsin (Thermo Scientific) was added, and tryptic digestion was performed under agitation at 37°C for 30 minutes, 2 hours, 4 hours, or 18 hours. Upon completion of each timepoint, samples were centrifuged at 13,000 rpm for 5 minutes to separate any undigested CDM material, and peptide-containing supernatants were transferred to a clean tube, acidified using 50% trifluoroacetic acid (TFA) and stored at -20 °C.

#### Time-lapsed tryptic digestion

For time-lapsed tryptic digestion, trypsin was added following the pre-digestion, and the tryptic digestion was continued under agitation at 37 °C (n = 2). After 30 minutes, samples were centrifuged at 13,000 rpm for 5 minutes, and the supernatants were transferred to a clean tube, while the remaining proteins were resuspended in 2 M urea in 100 mM NH_4_HCO_3_, and the digestion continued at 37 °C with addition of equal amount of trypsin. After 2 hours since the initial trypsin addition (or 1.5 hour after addition of the second batch of trypsin), samples were centrifuged, and the process was repeated, and the 4-hour and 18-hour timepoint samples were obtained. Samples were acidified with 50% TFA upon timepoint completion and stored at -20 °C until further processing.

Acidified peptides were desalted on C18 desalting columns (Thermo Scientific) following the manufacturer’s instructions and eluted in 50 % acetonitrile (ACN) solution containing 0.1 % TFA. Peptides were lyophilized for 4 hours and reconstituted in 5 % ACN solution containing 0.1 % formic acid. For all samples, peptide concentration was measured using a colorimetric peptide quantification assay (Thermo Scientific). Finally, the concentration of the samples was adjusted to 250 ng/μL.

### LC-MS/MS analysis

#### Data acquisition

1μg-equivalent of peptides from each sample was dissolved in 15uL 0.1% formic acid and injected and separated by a capillary C18 reverse-phase column of the mass-spectrometer-linked Ultimate 3000 HPLC system (Thermo Fisher) using a 90-minute gradient (5-85% acetonitrile in water) at a 300 nL/min flow rate. Mass spectrometry data were acquired on a Q Exactive HF mass spectrometry system (Thermo Fisher) with nanospray ESI source using data-dependent acquisition.

#### Database search

Mass spectrometry raw files were converted into .mzML files using msConvert [57], searched and filtered by MSFragger [58] (version 3.2) and Philosopher [59], respectively. The database search was performed against a custom proteome reference database generated by amending standard mouse reference proteome (UniProt Mus musculus C57BL/6J proteome last modified on March 7, 2021, and including 110,926 entries, 55,463 forward sequences and 55,463 decoy sequences) with human SNED1’s protein sequence (UniProt Q8TER0). Trypsin was specified as cleavage enzyme allowing 1 missed cleavage, mass tolerance was set to 50 ppm for precursor ions and 20 ppm for fragment ions. The search included variable modifications of methionine oxidation, lysine and proline hydroxylation [60], asparagine, and glutamine deamidation, glutamine to pyroglutamate, and fixed modification of carbamidomethyl cysteine. Peptide length was set within a range between 7 to 50 amino acids and false discovery rate (FDR) was set to 1% at the peptide/spectrum match (PSM), peptide, and protein levels.

### Data analysis

We restricted our analysis to reviewed UniProt entries and annotated the dataset to identify ECM and ECM-associated proteins according to the matrisome nomenclature [17,30]. For protein identification, we further restricted our analysis to proteins detected with at least 2 peptides. Proteome coverage was calculated from mapping all identified peptides to custom proteome reference sequences described above.

#### Semi-quantitative analysis of protein abundance

For each protein, the normalized spectral abundance factor (NSAF) was calculated as described by Zybailov et al. [42]. In brief, NSAF is calculated as the number of spectral counts (SC) identifying a protein, divided by the protein’s length (L), divided by the sum of SC/L for all proteins of the dataset [43].

#### Analysis of sequence coverage

To investigate the aggregated coverage at a given timepoint of the time-lapsed tryptic digestion, identified peptides from previous and current timepoints were pooled together prior to peptide mapping and coverage calculation (*e*.*g*., aggregated coverage at 18-hour was calculated based on all identified peptides from the sequential 30-minutes 2-hour, 4h-hour and 18-hour digestions). Change in sequence coverage when comparing the 18-hour stand digestion method and the data aggregated from the time-lapsed digestion method was calculated as follows:

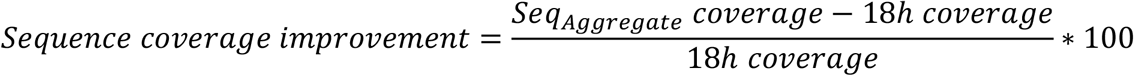

#### Dissimilarity index was calculated as follows

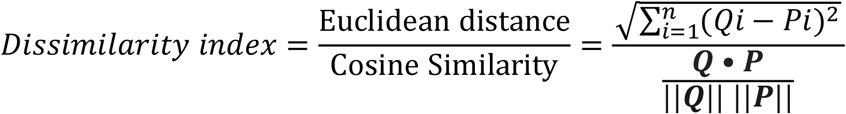

Where n = 4 (4 independent timepoints). Q and P represent the vectors consisting of 4 aggregated coverage in control or SNED1-overexpressing samples, respectively.

#### Statistical analysis

The data used for comparison between the coverages in proteins grouped by ECM categories was first checked for normality using the Shapiro-Wilk test [61]. Considering the non-normality of the data, the distribution of standard 18-hour digestion coverages and the aggregate coverage counterparts were compared using Wilcoxon signed-rank test, with p-values reported following Bonferroni adjustments for stringent results [62].

#### Data visualization

Proportional 2-circle Venn diagrams were generated using the Venn Diagram Generator: http://barc.wi.mit.edu/tools/venn/.

Visualization of the intersection of multiple datasets was performed using UpSetR [63] and the online application: https://gehlenborglab.shinyapps.io/upsetr/.

Visualization of the coverages and NSAF values was performed using Morpheus: https://software.broadinstitute.org/morpheus.

#### Peptide mapping on 1D representation of protein structures

The information of domain start and end positions were obtained from the Simple Modular Architecture Research Tool (SMART [64]). Each peptide was assigned the domains using a heuristic created in an Excel file that determined where the start and end position of the identified sequences belonged to (*see Supplementary File 1*). The outliers that were matched onto the non-domain regions of the protein were manually excluded in the count of spectra count ratio on the model of what has been described by Eckersley [65].

#### Peptide mapping on 3D representation of protein structures

The pdb files of AlphaFold prediction models were obtained for the proteins of interests [45]. Each pdb file was visualized and annotated using UCSF Chimera (https://www.cgl.ucsf.edu/chimera/) by selecting the sequences of interest to modify their colors manually onto the model.

## Data availability

All scripts were written in Python 3.8 and are available on the Gao lab website at http://pepchem.org:35091/blackjack/sequential_sned1/tree/master.

Raw mass spectrometry data have been deposited to the ProteomeXchange Consortium [66] via the PRIDE partner repository [67] with the dataset identifier PXD030713. *The raw data will be made publicly available upon acceptance of the manuscript*.

## CRediT AUTHOR STATEMENT

**Fred Lee:** Formal analysis, Investigation; Methodology; Visualization; Roles/Writing - original draft.

**Xinhao Shao:** Formal analysis, Investigation; Methodology; Software; Visualization; Roles/Writing - original draft.

**Yu Gao:** Conceptualization, Formal analysis, Funding acquisition, Investigation; Methodology; Resources; Software; Supervision; Validation; Roles/Writing - original draft.

**Alexandra Naba:** Conceptualization, Formal analysis, Funding acquisition, Investigation; Methodology; Project administration; Resources; Supervision; Validation; Roles/Writing - original draft.

## ACKNOWLEDGMENTS

The authors would like to thank all the members of the Naba and Gao labs for their feedback on the project.

## FUNDING SOURCES

Funding for this project was also provided in part by the University of Illinois Cancer Center at Chicago through the UICC Pilot Project funds (award 2020-PP-07 to YG and AN). This work was also supported in part by a start-up fund from the Department of Physiology and Biophysics, a pilot grant from the Cancer Biology Program at the of Illinois Cancer Center at Chicago, and an NIH R01 (R01AR074997) to AN and by an NIH R35 (R35GM133416) MIRA award to YG.

## CONFLICT OF INTEREST

AN has a sponsored research agreement with Boehringer-Ingelheim for work unrelated to the research presented in this study. The other authors declare that they have no conflict of interest.

## SUPPLEMENTARY FIGURE LEGENDS

**Supplementary Figure 1. Time-lapsed tryptic digestion increases overall sequence coverage**

**A**. Each line of the line plots represents, for a given matrisome protein, the average sequence coverage calculated at each timepoint of the time-lapse digestion. Each graph represents the data for a given matrisome category.

**B**. Violin plot represents the distribution of average sequence coverage between the standard 18- hour digestion protocol (blue), the coverage obtained at the 2-hour timepoint of the time-lapsed digestion protocol (orange), and the coverage obtained by aggregating data from the 4 timepoints of the sequential digestion (grey). The median is represented by the dotted line and the interquartile range is indicated by the colored dotted lines. One-way ANOVA was performed to compare the groups, and the p-value was reported (*p<0.05, **p<0.01, ***p<0.001, ****p<0.0001).

**C**. To rule out that the increase in the sequence coverage percentages seen upon aggregation of data from the different digestion timepoints was due to the simple increase in the number of datasets collated, we aggregated four datasets obtained using the standard 18-hour digestion method (*i*.*e*., four files) and compared the average aggregated coverage (left violin plots) to the average aggregated coverage obtained by integrating data from the four distinct digestion timepoints (right violin plots). We confirm that the increased percentage of sequence coverage is indeed due to the benefit of the time-lapsed digestion method.

**Supplementary Figure 2. Time-lapsed tryptic digestion increases overall sequence coverage** Histograms represent the distribution of average coverage percentage of all ECM proteins at each timepoint of the time-lapsed trypsin digestion. Scatter plots compare the log2-transformed average sequence coverage of all identified matrisome proteins at different timepoints of the time-lapsed digestion. Proteins aligned along the diagonal indicate a similar coverage between the two methods.

**Supplementary Figure 3. Proteomic profiling of the composition of the ECM produced by SNED1_GFP-overexpressing MEFs**

**A**. Bar graph depicts, for each matrisome category, the number of matrisome proteins identified in two biological replicates (intersection), in CDMs produced by MEFs overexpressing SNED1_GFP, using different trypsin digestion durations either individually (left) or sequentially (right) (*see also Supplementary Table 4*).

**B**. Venn diagram shows the overlap between matrisome proteins identified at each timepoints of the sequential digestion. For each timepoint, the protein set is defined as the ensemble of proteins identified in two biological replicates.

**C**. Matrisome proteins identified at each of the four sequential timepoints or the standard 18-hour single timepoint digestions are compared, and the intersections are visualized using an UpSet plot (*see also Supplementary Tables 4 and 5*).

**D**. Venn diagram illustrates the comparison between matrisome proteins identified using a standard 18-hour digestion protocol or by aggregating data from the 4 sequential digestion timepoints of the time-lapsed protocol.

**E**. Each line illustrates, for a given matrisome protein, its cumulative average sequence coverage percentage at each timepoint of the time-lapsed digestion (*see also Supplementary Table 3*).

**F**. Violin plots compare, for each matrisome category, the distribution of average protein sequence coverage obtained with a standard 18-hour digestion protocol or by aggregating data from the 4 timepoints of the time-lapsed digestion. Median and interquartile range are indicated by the white dots and black bars, respectively. Wilcoxon signed-rank test was used to compare the distribution, and p-value was reported after Bonferroni adjustments were made (*p<0.05, **p<0.01, ***p<0.001, ****p<0.0001, ns: not significant).

**G**. Heatmap represents the normalized spectral abundance factor (NSAF) obtained for each matrisome protein produced by SNED1-overexpressing MEFs and detected using the time-lapse trypsin digestion protocol or the standard single timepoint (18-hour) digestion protocol.

**H**. Histogram represents the distribution of average coverage percentage of all matrisome proteins at each timepoint of the time-lapsed trypsin digestion detected in CDMs produced by SNED1-overexpressing MEFs. Scatter plots compare the log2-transformed average sequence coverage of all identified matrisome proteins at different timepoints of the time-lapsed digestion. Proteins aligned along the diagonal indicate a similar coverage between the two methods.

## SUPPLEMENTARY TABLES

**Supplementary Table 1**. Sample list and MS metrics

**Supplementary Table 2**. Proteomic profiling of cell-derived matrices produced by immortalized *Sned1*^*KO*^ mouse embryonic fibroblasts (MEFs) overexpressing GFP alone

**Supplementary Table 3**. Overview of matrisome proteins identified CDMs made by immortalized *Sned1*^*KO*^ mouse embryonic fibroblasts overexpressing GFP alone

**Supplementary Table 4**. Proteomic profiling of cell-derived matrices produced by immortalized *Sned1*^*KO*^ mouse embryonic fibroblasts overexpressing GFP-tagged SNED1

**Supplementary Table 5**. Overview of matrisome proteins identified CDMs made by immortalized *Sned1*^*KO*^ mouse embryonic fibroblasts overexpressing GFP-tagged SNED1

**Supplementary Table 6**. Comparison of matrisome proteins identified in CDMs lacking- and containing SNED1

**Supplementary Table 7**. Semi-quantitative analysis using normalized spectral abundance factor (NSAF)

**Supplementary Table 8**. Complete peptide report

## SUPPLEMENTARY FILES

**Supplementary File 1**. Example of a peptide mapping heuristic (.xlsx)

**Supplementary File 2**. Time-resolved peptide mapping on the 3-D AlphaFold model of SNED1. Video shows a full rotation around the y-axis (.mp4)

**Supplementary File 3**. Time-resolved mapping of peptides identified in control CDMs on the 3-D AlphaFold model of nidogen 2. Video shows a full rotation around the y-axis (.mp4)

**Supplementary File 4**. Time-resolved mapping of peptides identified in SNED1_GFP-containing CDMs on the 3-D AlphaFold model of nidogen 2. Video shows a full rotation around the y-axis (.mp4)

**Supplementary File 5**. Time-resolved mapping of peptides identified in control CDMs on the 3-D AlphaFold model of thrombospondin 1. Video shows a full rotation around the y-axis (.mp4)

**Supplementary File 6**. Time-resolved mapping of peptides identified in SNED1_GFP-containing CDMs on the 3-D AlphaFold model of thrombospondin 1. Video shows a full rotation around the y-axis (.mp4)

