## Supplementary Figures for "Time-lapsed proteomics reveals a role for the novel protein, SNED1, in modulating ECM composition and protein folding"

### Supplementary Figure 1

A.

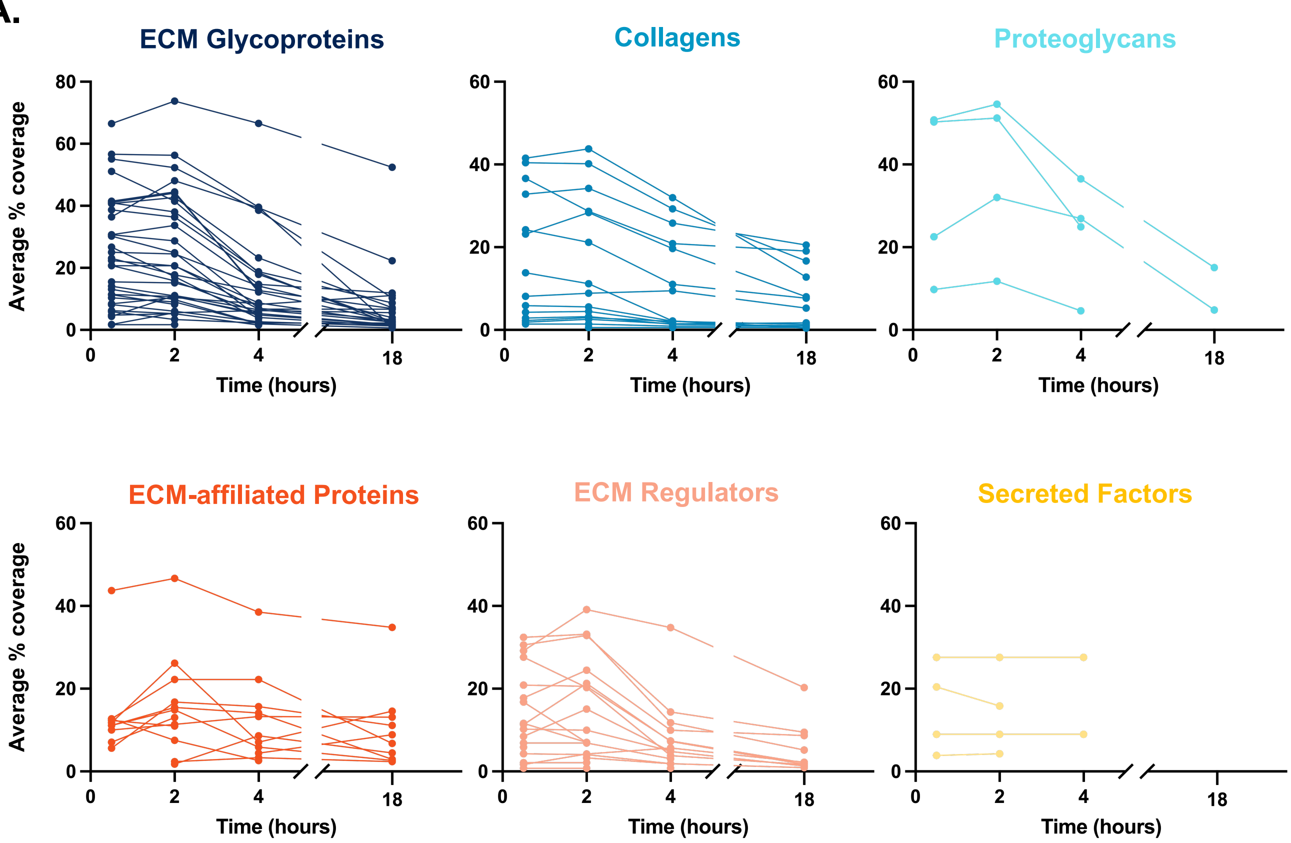


B.

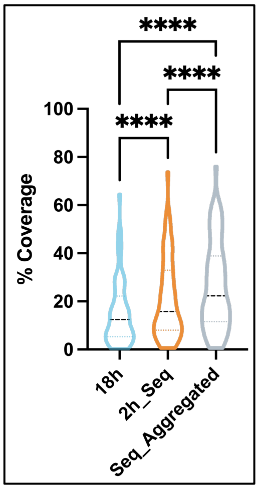


C.

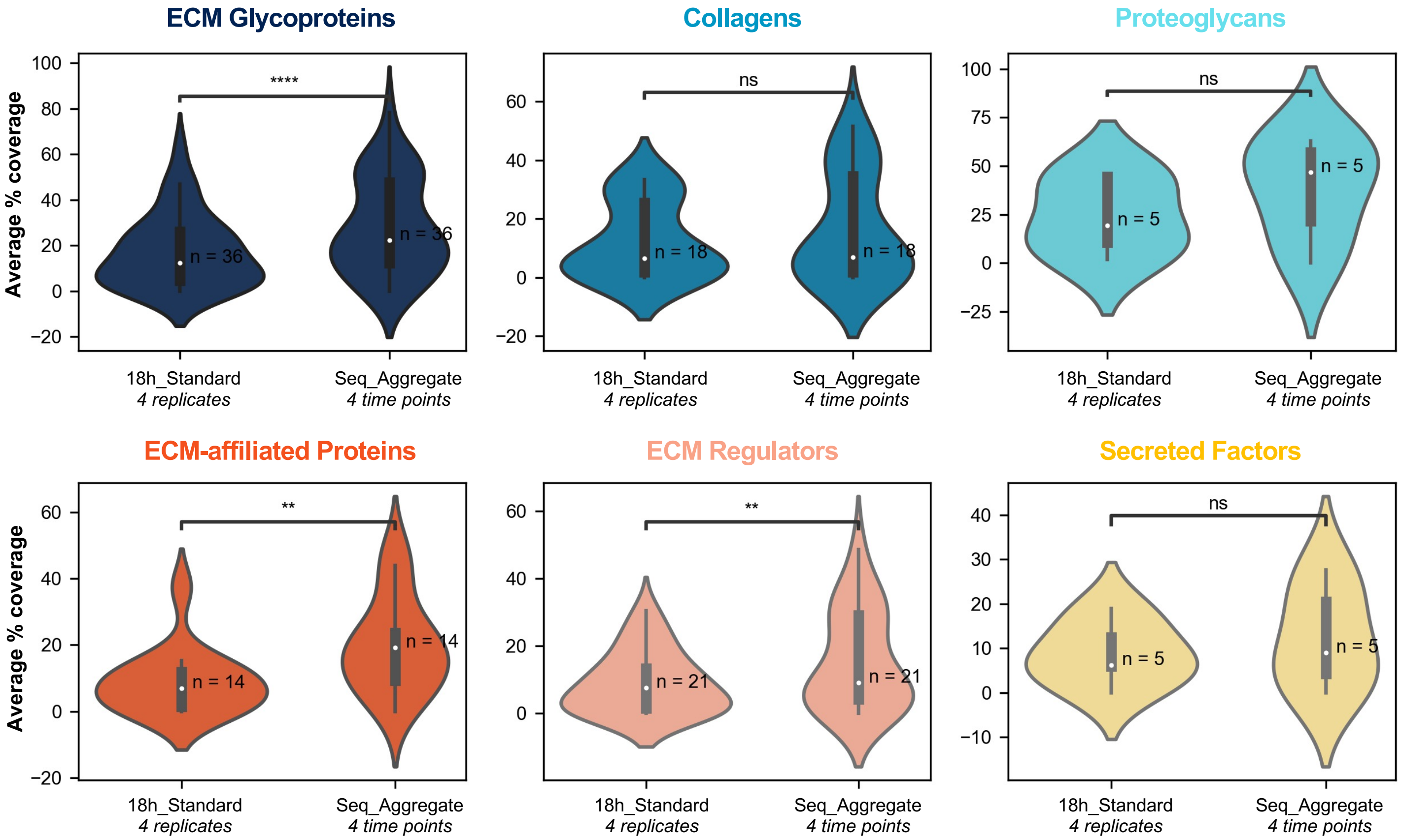

### Supplementary Figure 2

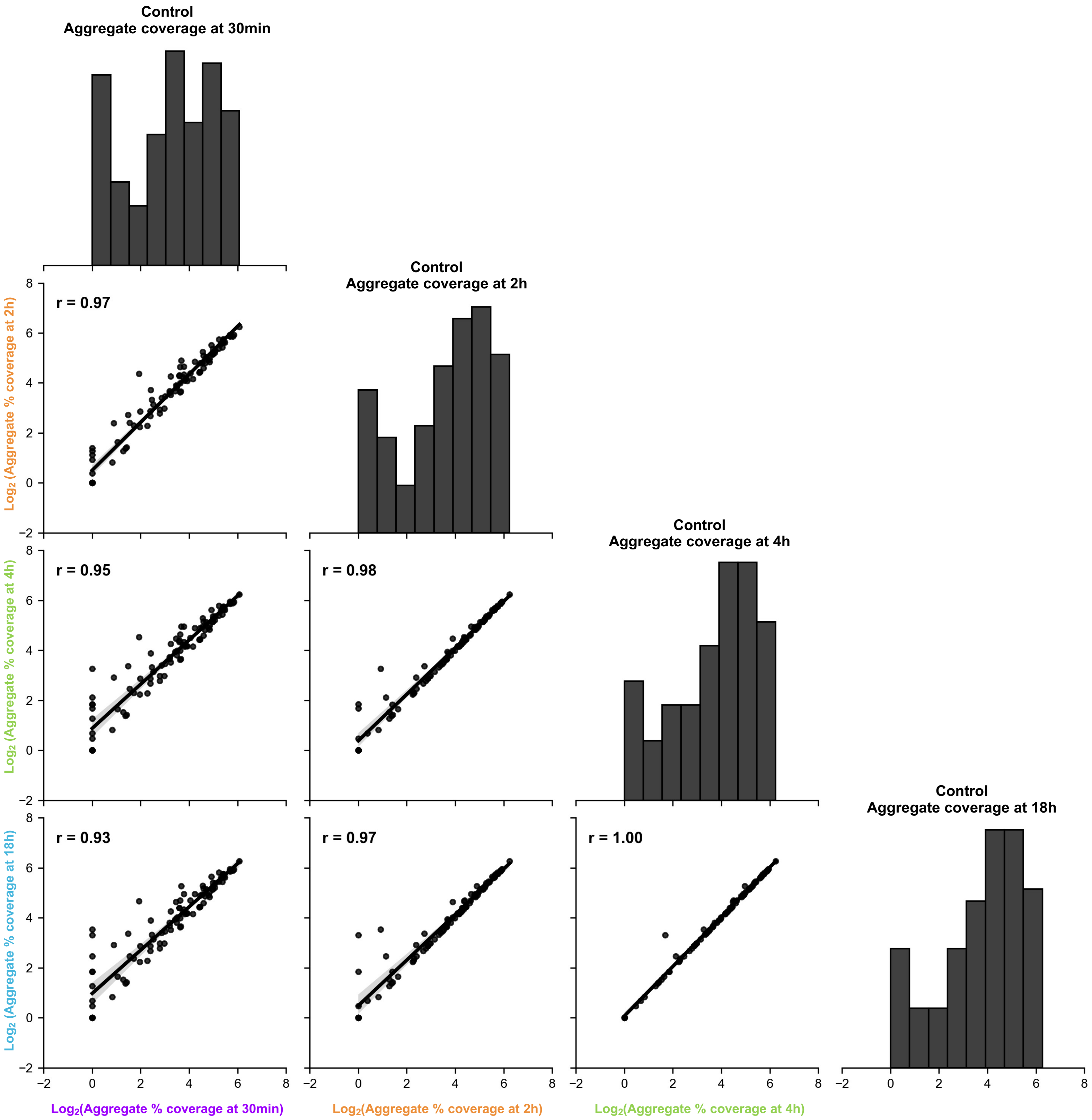

#### Supplementary Figure 3

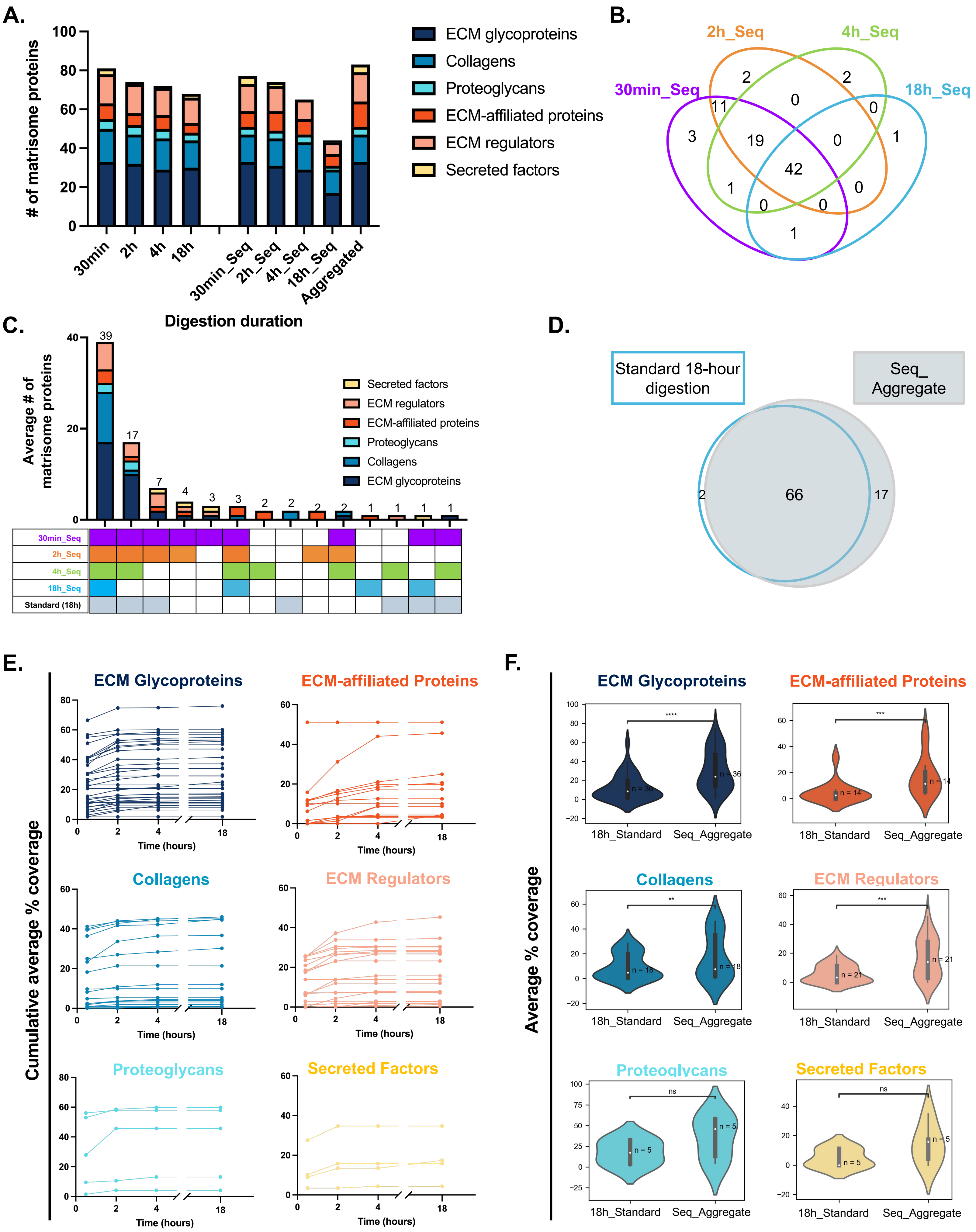

### Supplementary Figure 3

## G.

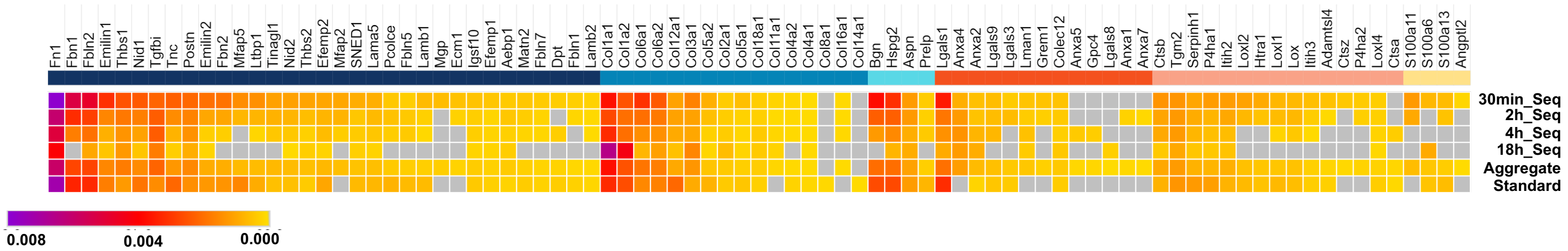


## H.

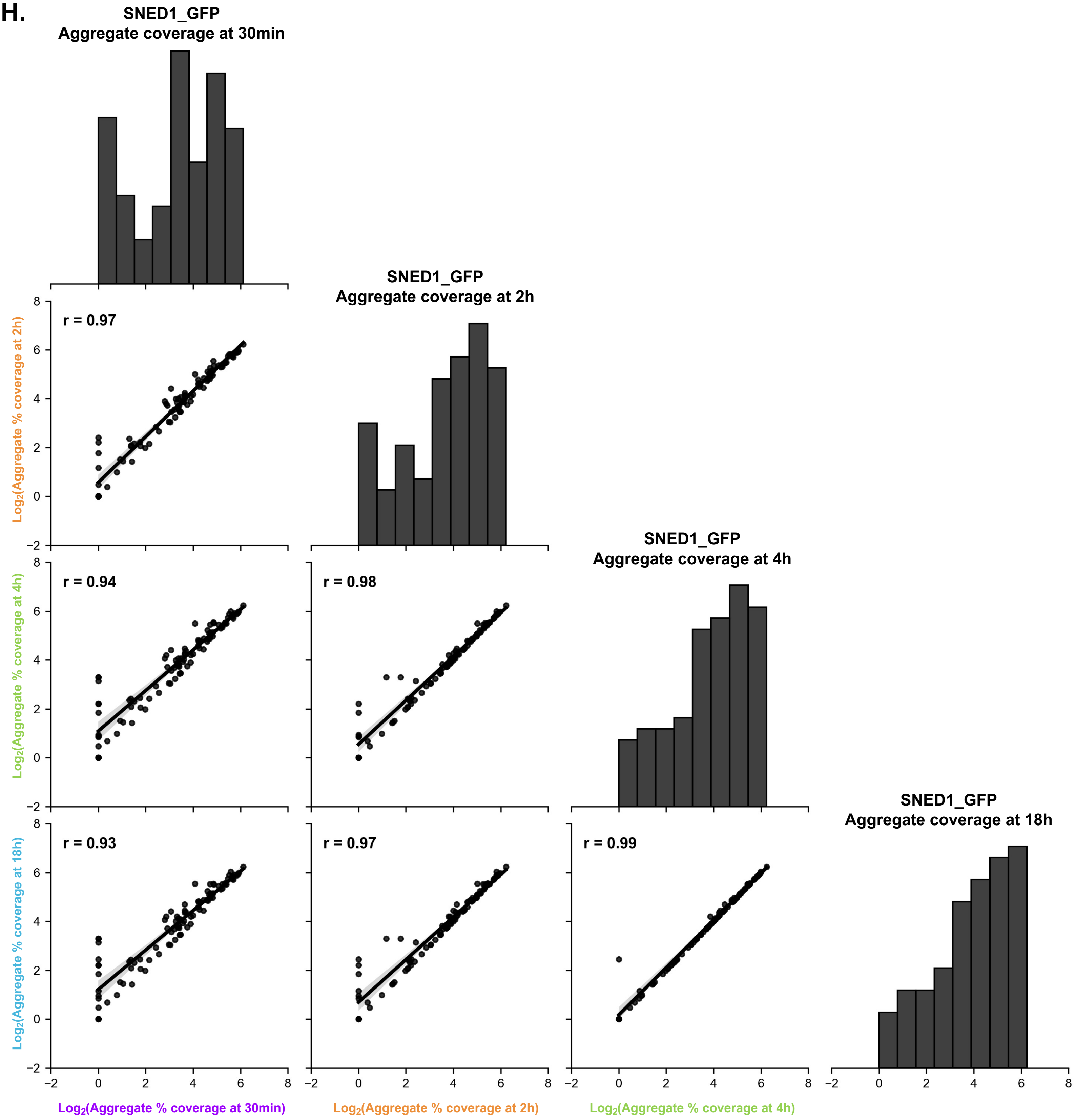
